## Supplementary Information for main manuscript for "Convergent Transcriptomic Evidence Reveals the Dysfunctional Quantitative Mechanism of Synaptic Plasticity Control in ASD"

**Supplementary Figure 18**

**Supplementary Table 30**

### Supplementary Note 1 The evidences for the endophenotype of synaptic plasticity

#### Synapse-related dysfunction feature by Gene enrichment analysis in ASD

##### Gene enrichment analysis

Gene Enrichment Analysis (GEA) is a widely used approach to identifying biological connections. We implemented the hypergeometric model to assess whether the number of selected genes in the ASD associated with interested items of biology knowledge is larger than that which can be expected purely by chance. In particular, the P-value determines whether any term annotates a specified list of genes at frequency greater than that which can be expected by chance, as determined by the hypergeometric distribution:

$$p = 1 - \sum_{i=0}^{k-1} \frac{\binom{N-M}{n-i}\binom{M}{i}}{\binom{N}{n}}, \tag{1}$$

where $N$ is the total number of genes in the background distribution, $M$ is the number of genes within that distribution that are annotated (either directly or indirectly) to the node of interest, $n$ is the size of the list of genes of interest, and $k$ is the number of genes within that list which are annotated to the node. The background distribution by default is all the genes that have an annotation. Then, the outcome of a statistical hypothesis test for the hypergeometric distribution in the gene enrichment analysis, divided by the total number $N$ of genes in the analysis, defines the enrichment efficiency $\eta$:

$$\eta = -\log_{10} P/N. \tag{2}$$

##### Progressive enrichment analysis in Reactome

We made a progressive enrichment anlysis from database of Reactome to further investegate the items related with DEGs. Synapse plasticity is a crucial endophenotype in ASD by the evidence from transcriptome enrichment of single-cell sequence data which has also discovered by former genetic studies through genetic risk genes of ASD.

The DEGs were applied to GEA in the Reactome database. At this time, we adopted a novel strategy to continuously execute the GEA procedure to find the most significant enrichment items about these DEGs across various neuro-types in ASD. As shown in Fig. 1, we noticed that the Neuronal System and the Chemical Synapses are related to DEGs. After investigating the subtype of these two parent items, we found that the NMDA and corresponding postsynaptic events (Fig. 1B) are more likely (ES=23.6% and -log10 P=14.8) to explain the dysregulation of ASD gene expression. Further inquiries into this NMDA-related item, we saw Post NMDA receptor (ES=25.0% and -log10 P=14.0) and Glutamate binding (ES=45.4% and -log10 P=10.3) in Fig. 1C.

##### Enrichment analysis in KEGG

Besides the Reactome database, we performed KEGG enrichment analysis for ASD DE mRNAs, taking into account the various biological processes. GEAs in Signal Transduction (Fig. 2) and Neuronal System (Fig. 3) items of KEGG databases illustrated synapse and its plasticity may be the main feature in several ASD cell types.

#### The relationship of layer in cerebral cotex and glutamate neuron

The cerebral cortex is the outer layer of the brain that is responsible for many of the brain's higher functions, such as perception, cognition, and movement. The cortex is divided into six main layers that are numbered from the surface of the brain (layer 1) to the border with the underlying white matter (layer 6). Glutamate is the primary excitatory neurotransmitter in the brain and plays a crucial role in many cortical processes, including synaptic plasticity, learning, and memory. Glutamate neurons are a subtype of cortical neurons that use glutamate as their primary neurotransmitter. These cells are found throughout the cortex, and their distribution varies across cortical layers. The relationship between cortical layers and glutamate neurons is complex and varies depending on the specific region of the cortex. In general, however, glutamate neurons tend to be more abundant in the deeper layers of the cortex (layers 4-6) compared to the more superficial layers (layers 1-3). The distribution of glutamate neurons across cortical layers is thought to reflect the functional specialization of different cortical regions. For example, regions of the cortex that are involved in sensory processing tend to have a high density of glutamate neurons in the middle layers (layer 4), while regions involved in motor function tend to have a high density of glutamate neurons in the deeper layers (layers 5-6). Overall, the relationship between cortical layers and glutamate neurons reflects the complex organization and functional specialization of the cerebral cortex.

##### The relationship of ASD and intermediate neuron

Autism Spectrum Disorder (ASD) is a neurodevelopmental disorder that affects communication, social interaction, and behavior. Intermediate neurons (INs), also known as interneurons, are a type of neuron found in the central nervous system that help to integrate and modulate signals between other neurons. While research on ASD is ongoing, there is currently no evidence to suggest a direct relationship between ASD and INs. However, studies have shown that alterations in the development and function of certain types of neurons, including INs, may be involved in the pathogenesis of ASD. For example, some studies have suggested that reduced numbers or altered functioning of certain types of interneurons may contribute to the cognitive and behavioral deficits observed in individuals with ASD. Other studies have suggested that alterations in the development and connectivity of excitatory neurons may also play a role in the development of ASD. Overall, while there is no direct relationship between ASD and INs, ongoing research is helping to shed lights on the complex neurobiological underpinnings of this disorder.

##### INs with LTP or LTD

INs, like other neurons in the central nervous system, have glutamate receptors such as AMPA and NMDA receptors. These receptors play a crucial role in mediating synaptic transmission and plasticity in the brain. Additionally, studies have shown that INs can exhibit both long-term potentiation (LTP) and long-term depression (LTD), which are forms of synaptic plasticity that underlie learning and memory. For example, research has shown that LTP can be induced in hippocampal interneurons, which are a subtype of IN, in response to high-frequency stimulation. Similarly, LTD can also be induced in interneurons, leading to a decrease in synaptic strength. Overall, the ability of INs to express both AMPA and NMDA receptors and exhibit synaptic plasticity suggests that they play an important role in modulating the activity of other neurons in the central nervous system and in contributing to various forms of learning and memory.

#### Supplementary Note 2 The pipeline of constructing the mSiReN

##### Initial nodes and final nodes in mSiReN

##### DEGs connected with translation control of synapse plasticity in ASD

Among the molecules listed, the part most strongly connected with glutaminergic neurons and ASD is the glutamate receptor family, which includes AMPA receptors, NMDA receptors, and metabotropic glutamate receptors (mGluRs). Glutamate is the primary excitatory neurotransmitter in the central nervous system, and abnormalities in glutamate signaling have been implicated in ASD.

Specifically, dysregulation or dysfunction of glutamate receptors, particularly the NMDA receptor subtype, has been associated with ASD. NMDA receptors are involved in synaptic plasticity, learning, and memory, and their proper functioning is crucial for normal brain development and function. Alterations in the expression or activity of NMDA receptors have been observed in individuals with ASD, suggesting a link between glutamate receptor dysfunction and the pathophysiology of ASD.

Furthermore, metabotropic glutamate receptors (mGluRs) play a role in modulating synaptic transmission and plasticity, and aberrant mGluR signaling has also been implicated in ASD. Certain genetic variations affecting mGluR signaling have been associated with an increased risk of ASD. some proteins or protein complexes that are known to be involved in synapse plasticity, long-term potentiation (LTP), and long-term depression (LTD):

All mRNA precursors of AMPA, which served as LTP, named GluAs, are upregulated. All the ARC precursors which served as LTD labelled as ARC or ARP are downregulated. The expressions of these DEGs show that ASD may relate to LTP, leading to the dysfunction of the translation signal network. For these proteins or protein complexes closely with translation control of synapese plasticity, we illustrated their expression characteristics in volcanic map, see Figure 6.

I. The PIK3 pathway is regulated by multiple factors. It can be activated through TRK receptors (TRK->PIK3CA/PIK3CB) and G protein-coupled receptors (GPCR, specifically mGlu-> PIK3CG). Additionally, the complex "PIK3CA_PIK3R1" is regulated by the IGF pathway (IGF1R) and the RAS pathway (HRAS/NRAS/KRAS). These regulatory relationships highlight the intricate control mechanisms governing the PIK3 pathway activation.

II. The names Class IA and Class IB are based on the PI3K catalytic subunits. Humans express three classes of PI3K catalytic subunits, where Class IA specifically refers to the I class catalytic subunits. Mammals express four Class IA catalytic subunits (p110$\alpha$, $\beta$, $\delta$ and $\gamma$) encoded by PIK3CA, PIK3CB, PIK3CD, and PIK3CG, respectively. They can all phosphorylate PIP2 into PIP3. The expression of p110$\alpha$ and p110$\beta$ proteins is widespread, while p110$\gamma$ and p110$\beta$ expression is enriched in immune cells. Among them, P110$\alpha$, $\beta$, $\delta$ (PIK3CA, PIK3CB, PIK3CD) belong to Class IA, and p110$\gamma$ (PIK3CG) belongs to Class IB. Therefore, studying Class A/B in the PI3K pathway is sufficient.

III. Metabotropic glutamate receptors (mGlu receptors) are a class of G protein-coupled receptors that play important roles in modulating synaptic transmission and neuronal activity. The composition of mGlu receptors includes the following subtypes: K04603 (GRM1), K04605 (GRM2), K04606 (GRM3), K04607 (GRM4), K04604 (GRM5), K04608 (GRM6), K04609 (GRM7) and K04610 (GRM8).

IV. "iGlu":Ionotropic glutamate receptors are a class of ligand-gated ion channels that mediate the majority of fast excitatory synaptic transmission in the central nervous system. The composition of ionotropic glutamate receptors includes the following subtypes: K05197 (GRIA1), K05198 (GRIA2), K05199 (GRIA3), K05200 (GRIA4), K05201 (GRIK1), K05202 (GRIK2), K05203 (GRIK3), K05204 (GRIK4), K05205 (GRIK5), K05206 (GRID1), K05207 (GRID2), K05208 (GRIN1), K05209 (GRIN2A), K05210 (GRIN2B), K05211 (GRIN2C), K05212 (GRIN2D), K05213 (GRIN3A) and K05214 (GRIN3B). Specifically GRIA1-4 and GRIN1, GRIN2A-D, GRIN3A-B, play a crucial role in synaptic transmission and neuronal signaling. The interaction between GRIN1 and GRIN2A/B/C/D forms the complex "GRIN1_GRIN2A/B/C/D," which is involved in mediating glutamate neurotransmission.

#### Match with annotation database

The steps to match original nodes PIK3, RAS, CAMK2 and the final node EIF4E to databases as follows:

- To find the interesting source nodes ("receptors and upstream signals"): PIK3R1, PIK3CG, PIK3CA, HRAS, NRAS, KRAS, RASA1, RRAS and CAMK2A, and target nodes ("translation control"): EIF4E and EIF4EBP1.
- Match the molecules in the post-translational modified network of the Omnipath database ("named import_post_translational_interactions") and search for pathways. Complexes are not necessary for selecting initial nodes: Although complexes are present in the initial signals, they do not need to be selected because only the initial signal nodes are chosen, and the signals will match the corresponding downstream nodes, including complexes, in the interaction network, which already includes information about these complexes.

Pathway annotation databases help determine which proteins and genes are involved in other interesting pathways within the network. If the nodes in the collected signal regulatory network match in the SignaLink pathway and SIGNOR databases, we can determine which genes correspond to the interested proteins and which genes do not participate in the signaling pathways. In the signal network obtained through literature mining, there are 44 "initial nodes", but after filtering by the signal network annotation, there are only 12 nodes remain. The latter can be further matched in molecular interaction databases to find regulatory relationships. see Table 4.

We listed nodes discarded and retained by the network during the matching process with subproject of Omnipath interaction database, named “import_post_translational_interactions”. The detailed informations sees Table 5.

Plus some super nodes which are not in the annotation network, then match the interaction database again. But finally, In addition to the neurotransmitter receptors, there also have three super nodes which can’t match any items in the interaction database, see Table 6.

#### Match with interaction database

We proceeded to match the molecular interaction databases. The databases “omnipath_interactions” and “pathwayextra_interactions” provided over 40,000 molecular interaction records, and their intersection yielded over 10,000 records. However, when searching for the maximum of eight pathways, there are over 1,000,000 records. To handle this, we extracted the nodes directly from the database network, matched them with the nodes from the literature-mined network, and then used the initial and terminal nodes of interest for further matching. This approach may result in a limited number of matched regulatory relationships or missing links since only a fraction of the thousands of regulatory relationships are considered when extracting over 100 nodes. Nevertheless, it remains a feasible solution.

If we were to match all nodes in the database initially and then search for subnetworks within the obtained global network, it would require significant computational resources. Moreover, there would be many molecular protein nodes appearing between the designated initial and final nodes in the signal network pathways, making further research challenging (additionally, after matching with manually collected network nodes, there are still over 600,000 nodes).

After matching the nodes from the literature-mined network with the “omnipath_interactions” and “pathwayextra_interactions” databases, we obtained a network of 159 relationships. Following analysis and discarding nodes based on specific conditions (curation_effort>=2 and consensus_direction==1), we found that the constructed mRNA and protein regulatory network lacked translation control proteins (FMRP and CYFIP1), neurotransmitter receptors (mGluR and NMDAR), and regulons (ADNP, EN2, P2A, NF1 and AMPA/STK11).

### Supplementary Note 3 The NIVaCaR Algorithm

#### Procedure Within Mathematical Representation

Assuming a regulatory network $G$ defined as a set of interactions $i$= 1,..., $n_r$ and a set of species (*i.e.* nodes) $j$ =1,...,$n_s$. Each interaction $i$ is an ordered pair of species of the form $S_i \rightarrow T^i$, where $S_i$, $T^i \in$1,...,$n_s$ are the source and target species respectively. The symbol details see **Tab. 7**. Moreover, the sign of $i$ is denoted with $\sigma_i \in$ {-1,1}, distinguishing between activations ($\sigma_i$ = 1) and inhibitions ($\sigma_i$ = -1). We also defined a set of cell types $k$ = 1,...,$n_e$. Where in each cell type a set of species are perturbed $I_{j,k} \in$ {-1,0,1} and a set of species are measured $c_{j,k} \in$ {-1,0,1}. Variables $x_{j,k} \in$ {-1,0,1} are introduced to denote the predicted activation state of species $j$ in cell type $k$.

We introduce variables $u^+_{i,k} \in$ {0,1} and $u^-_{i,k} \in$ {0,1}; $i$ = 1,...,$n_r$; $k$ =1,...,$n_e$ to denote the activity of interaction $i$ in cell type $k$.

The activation state of a interaction $i$ is defined by the activity of its source node $x_{S_i}$ and the interaction sign $\sigma_i$. The interaction has a potential to activate its target node (when $u^+_i$ = 1), if and only if $\sigma_i \cdot x_{S_i}$ = 1. This occurs in two cases: either the source node is activated ($x_j$ = $S_i$ = 1) and has an activating effect on its target node ($\sigma_i$ = 1); or the source node is inhibited ($x_j$ = $S_i$ = -1) and has an inhibiting effect ($\sigma_i$= -1). Vice versa, a interaction has the potential to downregulate its target node ($u^-_i$ = 1), if and only if $\sigma_i \cdot x_{S_i}$ = -1.

For a series of cell types k, If $u_+$ = 1 then interaction $i$ is active and can potentially up-regulate its target node; else if $u\_$ = 1 then interaction $i$ is active and can potentially down-regulate its target node. A interaction $i$: $S_i \rightarrow T^i$ is active and may up-regulate $T^i$ ($x^+_{i,k}$ =1), if $x_{j,k}$ = 1 and $\sigma_i$ = 1 or $x_{j,k}$ = -1 and $\sigma_i$ = -1; $j$ = $S_i$. On the other hand, a interaction $i$: $S_i \rightarrow T^i$ is active and may down-regulate $T^i$ ($x^-_{i,k}$ =1), if $x_{j,k}$ = 1 and $\sigma_i$ = -1 or $x_{j,k}$ = -1 and $\sigma_i$ = 1; $j$ = $S_i$.

The rules of interactions discussed above can be modelled as linear equality or inequality constraints as follows:

$$\begin{aligned} u^{+}_{i,k} &\geq \sigma_i x_{j,k}; i \in \{1, \ldots, n_{\mathrm{r}}\}; j = S_i; k = 1, \ldots, n_{\mathrm{e}} \\ u^{-}_{i,k} &\geq -\sigma_i x_j, k; i \in \{1, \ldots, n_{\mathrm{r}}\}; j = S_i; k = 1, \ldots, n_{\mathrm{e}} \end{aligned} \tag{3a}$$

$$u^{+}_{j,k} \leqslant 1 - u^{-}_{j,k};\ i \in \{1, \ldots, n_r\};\ k = 1, \ldots, n_e \tag{3b}$$

$$\begin{aligned} u^{+}_{j,k} &\leqslant \sigma_i x_{j,k} + u^{-}_{j,k};\ i \in \{1, \ldots, n_r\};\ j = S_i;\ k = 1, \ldots, n_e \\ u^{-}_{j,k} &\leqslant -\sigma_i x_{j,k} + u^{+}_{j,k};\ i \in \{1, \ldots, n_r\};\ j = S_i;\ k = 1, \ldots, n_e \end{aligned} \tag{3c}$$

Moreover, The variables $x^{+}_{j,k}$ ∈0,1 and $x^{-}_{j,k} \in \{0,1\}$ denote the potential of node $j$ being up (or down) regulated. node $j$ may be up-regulated ($x^{+}_{j,k} = 1$) if $\exists i$: $x^{+}_{i,k} = 1$ or $I_{j,k} = 1$. On the other hand a node may be down-regulated ($x^{-}_{j,k} = 1$) if $\exists$ $i$: $x^{-}_{i,k} = 1$ or $I_{j,k}$ = -1. The activation state that node $j$ will ultimately assume ($x_{j,k}$) is the sum of $x^{+}$ and $x^{-}$. Thus, if $x^{+}_{j,k} = 1$ and $x^{-}_{j,k} = 0$, then $x_{j,k} = 1$, else if $x^{+}_{j,k} = 0$ and $x^{-}_{j,k} = 1$, then $x_{j,k}$ = -1, else if $x^{+}_{j,k} = 1$ and $x^{-}_{j,k} = 1$, then $x_{j,k} = 1$, else $x_{j,k} = 0$.

The variables are determined during the linear programming optimisation according to the following set of constraints of causal reasoning principle.

$$\begin{aligned} x^{+}_{j,k} &\leq \sum_{i:T_i=j} u^{+}_{i,k}; i \in \{1, \ldots, n_{\mathrm{r}}\}; k = 1, \ldots, n_{\mathrm{e}} \\ x^{-}_{j,k} &\leq \sum_{i:T_i=j} u^{-}_{i,k}; i \in \{1, \ldots, n_{\mathrm{r}}\}; k = 1, \ldots, n_{\mathrm{e}} \end{aligned} \tag{4}$$

$$x_{j,k} = x^{+}_{j,k} - x^{-}_{j,k} + I_{j,k};\ j \in \{1, \ldots, n_s\};\ k = 1, \ldots, n_e \tag{5}$$

For the definition of the activation state of a node $x_{j,k}$, two cases can be distinguished: For the noninput nodes, the activity of these nodes is defined by the potentials of incoming reactions. If and only if at least one incoming reaction has the potential to activate ($u^{+}_{i} : T_{i=j} = 1$), the node can have the potential to be activated ($x^{+}_{j} = 0 \vee 1$). Vice versa, a node can only have the potential to be down-regulated ($x^{-}_{j} = 0 \vee 1$) if at least one incoming reaction has the potential to inhibit ($u^{-}_{i} : T_{i=j} = 1$). A node is then up-regulated ( $x_{j,k} = 1$), if there is exclusively a potential to be upregulated ($x^{+}_{j} = 1$), and is down-regulated ($x_j = -1$) if there is exclusively a potential to be downregulated ($x^{-}_{j} = 1$). If none or both of the potentials exist, the node will remain neutral ($x_j = 0$).

##### Removal of feedback loops from the signaling network

. In addition, feedback loops are removed given that effects mediated through those are highly dynamic and hardly interpretable from a static snapshot as in an interaction network. For instance, positive feedback loops can lead to internal signals independent to external perturbations and break the inference of pathway activities. These were constrained through a distance variable $d_{j,k}$. Thereby, the distance of all nodes connected to a perturbation node is set to a value larger than zero, while all others are defined to be zero.

For example, for node $j$ to be active ($x_{j,k} = 1$), it either has to be directly perturbed $I_{j,k} = 1$, or be activated by an upstream interaction $i$, such that $j = T^{i}$ and $x^{+}_{i,k}$ =1. However, if $n$ nodes form a positive cycle (a cycle where all interactions are positive), then one node will be able to activate the next all the way around the cycle, without the need for an external perturbation (or an incoming interaction transitively connected to a perturbation).

As a consequence, only nodes connected to a perturbation can be deregulated. The distance increases from source node to target node if the interaction is active, *i.e.* $u^{+}_{i} = 1 \vee u^{-}_{i} = 1$, and is not allowed to pass the distance threshold $M$, which is considerably larger than expected path lengths.

The variables $d_{j,k}$ 0 represents the distance of node $j$ from a perturbed node in cell type $k$. If node $j$ is not connected to a perturbed node, then $d_{j,k} = 0$, else $d_{j,k} > 0$. For node $j$ to be active, $d_{j,k} > 0$ has to hold true. If $d_{j,k} = 0$, then $x_{j,k} = 0$. The distance of node $j$ has to be greater than all of its upstream nodes at least by one (to enforce that the distance grows the further away from the input nodes we move), unless, the upstream interactions are not active ($i.e. u^{+}_{i,k} = x^{-}_{i,k}$ =0). Finally, the distance of any given node cannot be greater than the total number of interactions $M$ in the signaling network. The above may be formulated using linear constraints in the following manner:

$$x^{+}_{j,k} \leqslant d_j$$
$$x^{-}_{j,k} \leqslant d_j \tag{6a}$$

$$d_{Ti} \geq d_{Si} + 1 - M + x^{+}_{i,k} \cdot M$$
$$d_{Ti} \geq d_{Si} + 1 - M + x^{-}_{i,k} \cdot M \tag{6b}$$

$$d_j \leqslant M \tag{6c}$$

The above constraints prohibit the ILP algorithm from conserving a positive feedback loop in the solution and all the included interactions to be active, unless there is an input node in the loop. Assuming a loop like that is conserved, then the distance $d_{j,k}$ would increase indefinitely in the loop, making the ILP infeasible since $d_{j,k}$ is bound by $M$. Where $M$ is a sufficiently big number.

#### Using FC value of DEGs to discover the common core module by NIVaCaR

In the next step, we aimed to apply NIVaCaR to the translated mRNA interaction network derived from the protein signaling network. We want to use the DEGs as nodes to represent the activation or inhibition status of these gene nodes in ASD, relative to normal samples. We used the fold change (FC) values from Differential Expression Analysis (DEA) to determine this relative change. Using mSiReN as the prior knowledge network, we applied an integer linear programming model fitting using the differential expression values of the gene nodes. This helps us identify which nodes and pathways are activated in ASD and investigate whether there is specificity or commonality across different types of neurons.

At the algorithmic level, we select the DEGs for each cell type that show dysregulation in ASD at the RNA expression level. These genes represent the dysregulated response in ASD at the RNA level. The FC values of these genes were binarized to determine whether they are upregulated or downregulated in ASD cells. The framework consists of the following steps:

- DE gene expression: We start with gene expression data obtained from different scRNA-sequnce data. These data reflect the relative expression levels of genes across the cell types as well as CTL/ASD.
- Binaryzation: To simplify the analysis and focus on pathway activation, we binaryze the gene expression data. This involves setting a threshold to categorize genes as either ”active” or ”inactive” based on their expression levels. This step transforms the continuous gene expression data into a binary representation.
- NIVaCaR algorithm: We apply the NIVaCaR algorithm, which is based on causal reasoning and network identification, to the binary gene expression data. NIVaCaR aims to identify the activated pathways by analyzing the relationships between genes and network structure.
- Integer Linear Programming (ILP): To infer the most likely activated pathways, we utilize an ILP framework within the NIVaCaR algorithm. ILP allows us to optimize the pathway activation patterns based on the binary gene expression data.

The activated subnetworks illustrated in Fig. 4 for different excitatory cell types and Fig. 5 for inhibitory types from NIVaCaR results. After comparing the results of NIVaCaR with the gene expression, we found that EIF4EBP is not a DEGs (for example in L4, logFC=-0.0024, AveExpr=0.0216 and P.Value=0. 711894) but still follows a causal logic of an “activated signaling pathway.” This indicates that NIVaCaR identifies causal relationships in network regulation processes rather than solely relying on gene expression levels to determine potential dysregulated molecules (see Fig. 11).

### Supplementary Note 4 The ProComReN algorithm

The functional characteristics of eukaryotic cells are largely determined by the properties of their regulatory networks. Notwithstanding the vast amount of biological data accumulated over the past decades, a global model of the way these networks determine the phenotypes of both healthy and diseased cells remains elusive. One goal of systems biology is to understand these networks at the highest possible protein functional level, for example to devise therapeutic strategies. Mathematical modelling of regulatory networks allows for the discovery of knowledge at the system level. However, existing modelling tools are often computation-heavy (such as ode furmula) and do not offer intuitive ways (logical algebraic parameter, such as CNORfuzzy [1]) to explore the model, to test hypotheses or to interpret the results biologically.

Numerous mathematical approaches exist to optimize and train regulatory network models against steady-state experimental data. Of these, logical models [2] are of particular interest, as they are able to capture essential features of the system being modelled and generate biological insights, while requiring less prior knowledge and experimental observations than differential equation models [3]. In addition, logical models are in general more powerful than statistical models, as they incorporate the relational information embedded in the network structure, while statistical models aiming at reverse-engineering biological networks from high-throughput data implicitly consider all possible topologies [4]. Some successful applications include the logical models of yeast cellcycle protein network [5], gene regulatory networks [6], signalling networks [7].

In logical models of systems at steady-state, nodes represent the degree of activation of the constituents of the system at equilibrium and edges represent the logical functions between nodes. These functions can be either linear or non-linear functions of the parent nodes and are combinations of the fundamental "AND", "OR" and "NOT" Boolean functions. While Binary Boolean models [8] only consider full activation or complete absence, more quantitative approaches, for instance, Probabilistic Boolean Networks (PBNs) [9] and Dynamic Bayesian Networks (DBNs) [10] can account for intermediate or continuous activation values and allow the integration of data uncertainty. These approaches are usually analyzed by Monte Carlo approaches, which can be computationally demanding or non-intuitive to use.

Here, we have developed a computational approach ProComReN to efficiently contextualize logical models of regulatory networks (very suitable for signal transduction network) with biological measurements (sc-RNA sequence data) based on a probabilistic description (PBN) of rule-based interactions (or DBN with algebraic furmula) between the different species (molecules or nodes). This algorithm is inspired by FALCON [11], Fuzzy [3] and CNOprob [12].

The tool presents a computational approach to contextualize logical models of regulatory networks, specifically designed for signal transduction networks. It facilitates the integration of biological measurements, such as cell-specific sc-RNA sequence data, into the modeling process. The approach is based on a probabilistic description, utilizing either Probabilistic Boolean Networks (PBN) or Dynamic Bayesian Networks (DBN) with algebraic formulas, to capture the rule-based interactions between the different species or nodes within the network.

A Bayesian interpretation of the logical "gates" allows for an algebraic formulation of the system and an efficient calculation of the long-term steady-state of the system given the specified inputs. A gradient-descent optimizer is used to minimize the error function calculated as the sum of squared residuals between the simulated and experimentally measured nodes of interest. By employing this approach, a broader range of potential regulatory edges can be encompassed without the need to delve into the intricate molecular mechanisms underlying each individual interaction. This strategy effectively reduces the model parameters, enabling increased computational efficiency, and providing possibilities for training large-scale biological networks and facilitating the Post-hoc analyses detailedly.

It is important to note that the model operates on synchronized update data, as the translation control network is directly regulated by the signal transduction network. Subsequently, the model maps the translated information onto its mSiReN for analysis. Asynchronous updates are not considered in this context and the logical formulation of the interaction between different molecules is intuitive. ProComReN is a algorithm for the efficient contextualization of logical network models which provides important qualitative and quantitative information about the system being modelled. Specifically, ProComReN is well suited to assess the relative contributions of different signal regulatory mechanisms to the behavior of the system at steady-state.

#### The mathematical representation of the ProComReN algorithm

There is a clear conceptual difference between differential equation and coarse-scale models. The former can be used for a detailed representation of biochemical reactions, whereas the latter emphasize fundamental, generic principles between interacting components. In this context, the theory of computational models [10] classes that are both discrete-time and discrete-state are called coarse-scale models.

Limitations of differential equations for fine-scale modeling of biological interactions at the molecular level:

- Those models are computationally very demanding.
- The model selection problem is usually ignored
- The underlying biological system is assumed to be known.

So-called graphical models can overcome the above-mentioned modeling problems, and advanced analysis tools have been developed for them.

The use of holistic, coarse-scale models is also supported by the fact that the currently available data is limited both in quality and the number of samples. That is, there is no advantage using models that are much more accurate than the available data. Another constraint to be kept in mind is that the modeling framework should be selected on the basis of the preferred goals, i.e., to what kinds of questions are we seeking answers.

#### The relationship between PBN and DBN

Probabilistic Boolean Network (PBN) is a specific type of dynamic Bayesian network (DBN) that is used to model and analyze the behavior of complex systems, particularly in the field of computational biology. The main idea behind PBNs is to represent the state of each variable in the system as a Boolean value (either true or false) and introduce probabilistic transitions between the states of these variables over time. Unlike traditional Boolean networks, where the transitions between states are deterministic, PBNs incorporate uncertainty by assigning probabilities to the state transitions.

In a PBN, the system is modeled as a directed acyclic graph (DAG) consisting of Boolean variables as nodes and probabilistic dependencies as edges. Each node represents a Boolean variable, and the edges represent the probabilistic influence of the parent nodes on the state of the child node. The probabilities associated with the edges define the transition probabilities between the states of the variables. The transition probabilities in PBNs can be specified in different ways. One common approach is to use a Boolean function or a logical rule to determine the probability of transitioning from one state to another. These functions can be defined based on prior knowledge, experimental data, or expert opinions. PBNs are particularly useful for modeling and analyzing biological systems, such as gene regulatory networks. They allow researchers to capture the stochastic behavior and inherent uncertainties in biological processes. By simulating the dynamics of the PBN, researchers can gain insights into the behavior of the system, make predictions, and study the effects of perturbations or interventions.

The relationship between PBNs and DBNs lies in the fact that PBNs are a specific type of DBN. DBNs, as mentioned earlier, are a general class of models that capture temporal dependencies and uncertainties in dynamic systems. PBNs, being a specific instance of DBNs, focus on representing Boolean variables with probabilistic transitions. Therefore, PBNs can be seen as a specialized form of DBNs tailored for modeling Boolean systems with probabilistic behavior. It's worth noting that while PBNs are widely used in computational biology, DBNs have broader applications beyond Boolean systems and can model various types of variables with different probability distributions, continuous or discrete.

Boolean functions or logical rules play a crucial role in specifying the transition probabilities within PBNs. in a PBN, Boolean functions or logical rules are used to determine transition probabilities. In a PBN, each node can exist in different states, typically represented as binary values (e.g., "On" or "Off", "1" or "0"). The transition probabilities describe the likelihood or probability of a node transitioning from one state to another.

#### Explanation for edges in ProComReN

The passage describes the meaning and constraints of the weights associated with edges and hyperedges in a modeling framework. Each edge or hyperedge in the network is assigned a weight, denoted as $k_j^{(i)}$, which represents the relative influence of the upstream node to the downstream node. In this Bayesian-based modeling framework, the weights must adhere to the law of total probability. Specifically, for each node$X^{(i)}$ that has a set of $m$ activating functions denoted as $j_+$, the sum of the activating weights $\sum_{j_+=1}^{m} k_{j_+}^{(i)}$ must equal 1. This means that the weights associated with activating interactions should collectively account for the total influence on the downstream node. Similarly, for nodes that have a set of l inhibiting functions denoted as $j\_$, the sum of the inhibiting weights $\sum_{j\_=1}^{l} k_{j\_}^{(i)}$ must be between 0 and 1. This ensures that the weights of inhibiting interactions represent the relative inhibition of upstream nodes and are within a valid range. In summary, the weights in the modeling framework follow the principles of total probability, ensuring that the activating weights sum up to 1 and the inhibiting weights fall within the range of 0 to 1.

Given a network structure established from prior knowledge, a set of parameters (weights) and a set of experimental conditions, the steady-state of the network is computed for each of the conditions and the values of the nodes corresponding to the measured species are recorded. For each one of the conditions, the nodes of the network are initialized with random values, except for the nodes considered as inputs (external to the system) for which the value is determined by the experimental condition and kept constant. The network is then updated repeatedly by computing synchronously for each node the expected value of its probability distribution, given the value of its parent nodes and the weights associated with each interaction.

Because all nodes at each update are considered as independent, the inputs values of "AND" logical gates are multiplied. The computation of "OR" gates follows De Morgan's law, *i.e.* the complement of the union of two sets is the same as the intersection of their complements. Inputs pointing to the same child node that are not members of a logical gate are summed. Table 9 summarizes the different types of interactions explicitly formulated in our framework. The algebraic formulas used for the computations can be directly derived from the conditional probability tables of the DBN formulation of the logical interactions. The resulting dynamical system converges to a steady-state where each node value corresponds to the normalized equilibrium concentration of the activated form of the molecule in the system.

#### Assignment of additive effects to logic gates

There is one major difference on the ground assumptions should be discussed. When there is more than one interaction coming to a node, the CellNOpt packages assume that these interactions will either take the OR or the AND gate. Alternatively, ProComReN considers the third case like CNOprob i.e. an additive effect without assigning the OR/AND gate in this scenario. This assignment gives more flexibility to the ProComReN framework as it could be assigned more types of reactions with a broader set of transfer functions.

#### PBNs and Boolean functions or logical rules

The relationship between Probabilistic Boolean Networks (PBNs) and Boolean functions or logical rules lies in the specification of transition probabilities within PBNs. In PBNs, the transition probabilities represent the likelihood of transitioning from one state to another in the network. These probabilities can be determined using various approaches, and one common method is to use Boolean functions or logical rules. Boolean functions or logical rules provide a formal representation of the relationships and interactions among the variables (or nodes) in the network. These functions define the conditions under which a transition from one state to another occurs and assign probabilities accordingly. The Boolean functions can be derived from prior knowledge, experimental data, or expert opinions regarding the system being modeled.

By employing Boolean functions or logical rules, PBNs allow for a probabilistic description of the network dynamics. The transition probabilities are determined based on the satisfaction of the Boolean functions or logical rules, enabling the modeling of complex behaviors and capturing uncertainty in the system. In summary, Boolean functions or logical rules play a crucial role in specifying the transition probabilities within PBNs, facilitating the probabilistic modeling of regulatory networks and capturing the dynamics of the system.

#### A toy example for PBN and boolean functions or logical rules

a simple toy example to illustrate the relationship between Probabilistic Boolean Networks (PBNs) and Boolean functions. Imagine we have a regulatory network consisting of three genes: Gene A, Gene B, and Gene C. Each gene can exist in two states: 'On' or 'Off.' The behavior of the network is governed by the interactions between these genes. In a PBN, we assign transition probabilities to determine the likelihood of transitioning from one state to another. These probabilities can be determined using Boolean functions or logical rules. For our example, let's assume the following Boolean functions for each gene:

- Gene A: If Gene B is 'On' and Gene C is 'Off,' the probability of Gene A transitioning from 'Off' to 'On' is 0.8. Otherwise, the probability is 0.2.
- Gene B: If Gene A is 'On,' the probability of Gene B transitioning from 'Off' to 'On' is 0.6. Otherwise, the probability is 0.4.
- Gene C: If Gene A is 'On' and Gene B is 'On,' the probability of Gene C transitioning from 'Off' to 'On' is 0.9. Otherwise, the probability is 0.1.

These Boolean functions specify the conditions under which a transition occurs and assign corresponding probabilities. They can be derived from prior knowledge, experimental data, or expert opinions about the regulatory interactions between the genes. Using these Boolean functions, we can construct the transition probabilities for the PBN. These probabilities capture the dynamics of the network and allow for probabilistic reasoning about the system's behavior. For instance, based on the Boolean functions, if Gene A is currently 'Off' and Gene B is 'On,' the probability of Gene A transitioning to 'On' in the next time step is 0.2. Similarly, the probabilities of Gene B and Gene C transitioning between states can be calculated based on their respective Boolean functions and the current states of the genes. By simulating the PBN over time or performing probabilistic analyses, we can gain insights into the behavior of the regulatory network and the probabilities associated with different gene states. In summary, in a PBN, Boolean functions or logical rules are used to determine transition probabilities, which describe the likelihood of transitioning between states in the network. These functions capture the regulatory interactions and can be used to model and analyze the behavior of the network in a probabilistic manner.

#### Transitioning from one state to another

"State Transition" in the context of PBNs refers to the change in the state of a node or variable within the network. In a PBN, each node can exist in different states, typically represented as binary values (e.g., 'On' or 'Off', '1' or '0'). The transition probabilities describe the likelihood or probability of a node transitioning from one state to another. For example, let's consider a gene in a regulatory network represented by a PBN. The gene can be in an 'On' state or an 'Off' state. The transition probabilities associated with this gene would indicate the likelihood of it transitioning from an 'Off' state to an 'On' state or vice versa. These transition probabilities capture the dynamics of the network and provide information about the probability of a node changing its state based on the current states of its input nodes or other factors. They reflect the influence of the regulatory interactions and external factors on the behavior of the network. By specifying and analyzing these transition probabilities, we can understand how the network evolves over time and make probabilistic predictions about the states of the nodes in the network at different time points.

#### The segments of ProComReN algorithm

##### Objective function

To contextualize the model with experimental data, we employ a steady-state analysis approach. We extract the values of nodes in the network that correspond to the measured data and compare them with the normalized values obtained from experimental observations. The mean squared error (MSE) is then computed to quantify the discrepancy between the estimated values and the actual measurements. To optimize the weights and minimize this error measure, we utilize a gradient-descent algorithm. To ensure computational efficiency while accommodating varying degrees of recurrence in the networks, we employ the interior-point method [13]. This method allows us to strike a balance between accuracy and computational feasibility.

##### Differential regulation

In many real-life modelling applications, a system is studied in different contexts. For example, during a drug screen, the same signalling pathways are studied for different cell lines, or over time. One goal of systems biology is to identify differences between the contexts in the way the system is regulated. The same prior knowledge model is contextualized in parallel with different datasets corresponding to different contexts. ProComReN automates such analyses by optimizing identical models in parallel for multiple series of experimental conditions (cell types). We can discover which parts of the network are activated or shut down between cell lines/time points, and this may lead to the identification of specific interventions strategies for each context.

###### Rapid optimization

Using the gradient-descent optimization algorithm fmincon with interior-point method, ProComReN is able to rapidly estimate the set of weights that minimizes the objective function. Random initialization of the weights is done either from a uniform distribution across the [0, 1] range, or from a truncated normal distribution centred on 0.5.

###### Subsequent analyses on optimized logical networks

ProComReN implements post-optimisation analyses including systematic edge-knockout and systematic node-knockout. Based on the results from post-hoc analyses, we could obtain the list of nodes and edges which are sensitive and insensitive to perturbations. Those which are insensitive to node and edge knockouts could potentially be removed from the model. In contrast, those which are sensitive, i.e. Akaike Information Criterion (AIC) value increases substantially after the edge/node removal, implies that the respective component of the model is vital for model fitting and could potentially be the key regulatory point. Once a set of parameters has been inferred from a given topology and dataset, a series of additional analyses can be performed to gain more insight into the systems-level properties of the regulatory network being modelled. Subsequent analysis can identify context-specific parametrizations and topologies.

**Interactions knockouts.** ProComReN allows the systematic removal of each edge in the network and provides a graphical output showing the effect on the global fitness of the model. The models are compared using the Akaike Information Criterion (AIC) [14], which balances goodness-of-fit with model complexity. By using this additional analysis, it is possible to differentiate the crucial edges of the system from the ones that are dispensable, which can be pruned out.

**Nodes knockouts.** A frequent goal of systems biology analyses is to identify the crucial molecules of a regulatory network. Often performed via network topological properties (centrality measures), this identification is of particular interest in the case of target discovery efforts. ProComReN allows the systematic evaluation of models in which each node is removed from the network. The comparison of these models using the AIC allows to identify these crucial nodes not only from topological properties but from the effect their removal has on the behaviour of the entire system .

Node values, being comprised in the interval [0, 1], represent the probabilities for molecules to be in their active state at equilibrium. They can be understood as the normalized average activities of the nodes. The computed parameters, or weights, also comprised in the interval [0, 1] and subject to the law of total probability, represent the probabilities for the designated interactions to influence downstream nodes. They can be interpreted as the relative influences of the parent nodes on their children nodes and are useful in assessing the flow of the signal transduction.

#### Supplementary Note 5 The pipeline of mSiReN reduction for ProComReN algorithm

The criterion of filtering nodes in mSiReN:

1. FC value (P-value)
2. Expression isn't too low
3. Topology of network

4. Priori knowledge

To determine the checkpoint nodes in the upstream, two criteria need to be met: criterion 1, which applies to both some other nodes and upstream nodes, and criterion 2, which applies only to the other nodes. However, for the upstream nodes, they should meet criterion 2 but not criterion 1. This is because the control or ASD states refer to the entire system, including all nodes and edges, whereas the upstream nodes serve as a classification reference for other nodes, providing a binary criterion based on their expression. On the other hand, the downstream nodes should not meet criterion 2. Not only do these nodes need to pass the filter of 7 nodes and 10 edges in the inverse trace, see Fig. 12, but there are a lot of nodes that are crucial for the network but exhibit lower expression.

The significant difference between the method of the union network derived from the NIVaCaR results of cell types and the core module-based inverse trace network (Fig. 13) lies in the expression or fold change (FC) value of nodes in the network. In the former approach, NIVaCaR utilizes the FC values to filter and label nodes as negative or active, executing the algorithm to identify logically activated pathways or subnetworks. However, in the latter approach, nodes in the entire network are not selected based on FC values, and the ProComReN algorithm remains valid even if some nodes are not differentially expressed genes (DEGs) in the network. Therefore, if the ProComReN method is to be used, the original network should be employed instead of the network derived from the NIVaCaR results which includes filtered nodes based on FC values.

If nodes with expression lower than 20% in the specific cell type are deleted, the node EIF4EBP would be removed (Fig. 14). This general expression filter applied to the "translation control" part results in the deletion of EIF4EBP. However, the ProComReN analysis still handle this node appropriately. Despite its low expression, EIF4EBP can be included in the analysis, its input edges may be less activated in the network. The analysis can proceed even with one or two nodes exhibiting low expression levels.

The network represents a regulatory system with multiple nodes interconnected by edges. The upstream nodes including CAMK2A, HRAS, KRAS, NF1, PIK3CA, PPP2CB, PRKAA1, PRKAA2, PTEN, RAC1, STK11, and SYNGAP1 play crucial roles in influencing the downstream behavior of the network. Among these upstream nodes, CAMK2A and SYNGAP1 are isolated, indicating that they do not receive direct inputs from other nodes within the network. This isolation suggests that their regulatory effects may be unique or independent of other nodes.

Additionally, the network (Fig. 15) contains nodes such as MAP3K7 and MAP3K2, which are not considered part of the upstream nodes but still serve as input nodes based on the network's topology. These nodes may contribute to the overall behavior of the network, although they are not directly involved in the regulation of the upstream nodes. Certain nodes in the network, namely HRAS, PIK3CA, RAC1, PRKAA1, and PRKAA2, have incoming edges that are deemed important and should not be deleted. These edges likely represent significant regulatory interactions that are crucial for the proper functioning of the network. But based on the input nodes selecting rule: "Target more upstream pathway". Discard PPP2CB and PTEN.

Finally, we noticed some nodes have homolog such as AKT2, MAP2K1 and PRKAA1 (see Table 8), but it has lower expression in ASD. After filtering homolog with lower expression, AKT2, MAP2K2 and PRKAA1 (Fig. 16), we obtained the final netwrok structure (Fig. 17) which have 25 nodes and 43 edges containing RHEB, TSC2, RAF1, EIF4EBP1, MTOR, MAP2K1, NF1, MAPK1, MAP2K2, HRAS, KRAS, PTEN, MKNK1, PDPK1, AKT3, TSC1, PRKAA1RPS6KA5, AKT2, GSK3B, PRKAA2, STK11, PIK3CA, RAC1, PPP2CB, RPTOR, RPS6KB1 and EIF4E.

### Supplementary Note 6 Contextualizing mSiReN with ProComReN

#### Contextualizing mSiReN into the scRNA sequence data

Our capacity to generate large datasets is increasing steadily. A useful way to extract mechanistic insight from the data is by integrating them with a prior knowledge network of signalling to obtain dynamic models. Logic networks are among the conceptually simplest modelling frameworks. They capture the mechanistic relationship between molecular entities by logic gates. Due to their simplicity, they are highly scalable and widely applied. We hava constructed a general RNA network of protein (or complexes) precursors, mSiReN, which was further analyzed using the NIVaCaR algorithm. NIVaCaR have revealed the activity subnetworks for every cell types using the FC value of DEG nodes. This algorithm allowed us to identify the core downstream modular components of the network.

To understand the origin of these downstream components especially core module EIF4EBP1 and EIF4E, we have performed reverse tracing Fig. 12 to determine the sources of their regulatory signals, leading us to an interested general network. The molecular changes induced by perturbations such as drugs and ligands are highly informative of the intracellular wiring. For the detailed and coherent experimental data-based quantitative models, we contextualize mSiReN into sc-RNA sequence information directly.

#### Establishing the experimental data-based quantitative model

To gain a deeper understanding of the underlying regulatory mechanisms, we employed a boolean logic model, which allowed us to contextualize the gene expression data into a quantitative model. This model integrated the boolean logic rules with the experimental measurements, enabling us to capture the complex interactions within the regulatory network. In order to capture the cell-specific behavior, we employed an innovative method approach, which enabled us to create cell-specific quantitative models. These models incorporated the regulatory probabilities specific to each cell type, providing insights into the dynamics and behavior of the system at a cellular level. Hence, we established another pipeline to construct a Boolean logic model from a PKN and perturbation expression by ProComReN (inspired by the method of discrete logic modelling [7]) which integrates sc-RNA sequence data and puts the regulation network into a boolean logic model.

##### Experiment-liked data to traing model

We classified the high or low expression of a specific node in all cells belonging one neouron type to set the perturbation experiment data. Based on this node's (also gene) exprssion, we can continue to add another perturbation node. Those combinations of perturbation expression of nodes are similar with the data from the cell line experiment.

To optimize the model parameters and infer the regulatory relationships, we employed a combination of Genetic Algorithm (GA) and Integer Linear Programming (ILP) techniques. This optimization approach enabled us to refine the model and uncover the key regulatory elements associated with ASD. By applying the GA/ILP-based optimization, we identified the specific regulatory mechanisms that distinguish ASD from control samples. This approach provided valuable insights into the dysregulated pathways and potential molecular targets underlying ASD pathogenesis.

##### The preprocessing of data for the ProComReN model

Linear function normalization is a technique used to transform raw data into a range of [0-1], which simplifies downstream analysis while preserving the relationships present in the original data. This method assumes that the maximum and minimum values across the samples are similar. The formula for linear function normalization is

$$\frac{x - \min(x)}{\max(x) - \min(x)}, \tag{7}$$

where x represents the data points.

##### Use the sc-RNA expression data of CTL and ASD as the input of modeling

The proposed approach integrates the overall original mSiReN with single-cell RNA sequencing (scRNA-seq) expression data to achieve a comprehensive analysis of pathway networks. For detail, the mSiReN captures the intricate regulatory relationships among precursor mRNAs of proteins, while scRNA-seq expression data provides high-resolution information on gene expression patterns at the single-cell level. This combination of netwrok structure and information allows for a deeper understanding of the underlying regulatory mechanisms governing signal transduction network, facilitating the identification of key regulators and elucidating complex translation control process in ASD.

Therefore, we used the original whole mSiReN from interaction database through manual signal transduction network to execute the ProComReN directly (using gene expression) to establish a “quantitative” model.

##### Assess the dispency between two main cell types

Assessing the dispency of excitatory (L, L23, L4, L56 and L56CC) and inhibitory (IN, INPV, INSST, INSV2C and INVIP) ensured the network stability after the expression-based filter in the PKNs of different cell types for ProComReN. We calculated the variation of Standard Error (SE) for of expression for L and IN cell types respectively, **see Supplementary File 8**. The results show that L and IN are different majority type of cells based on interested nodes expression. So we therewith separated the main two cell types to ulteriorly analysis. On the other hand, more than 100 cells of input nodes remain to calculate in later steps after filter non-expression part Fig. 21. Input nodes in the main cell type ”Neu” (“Neumat”, “NeuNRGNI”, “NeuNRGNII”) are most zero expression. Deleting these cells will results in the unstable outcome of training ProComReN Fig. 22.

#### Supplementary Note 7 Relationship between NIVaCaR and ProComReN

##### General network structure and specific modeling

The constructed signal network is from literature and database, each cell type has the same signal network. Similarly, the network structure of mSiReN is for all cell types. However, for a specific cell type, the expressions of nodes and FC values are different, therefore the activated subnetworks and sub-pathways are unique in the followed modeling mothods.

With regard to NIVaCaR algorithm, each cell’s expression data (combining all cells within that cell type) is treated as an independent unit. The collected and organized signal network, together with the signal network built using Omnipath, is used as the target network for performing NIVaCaR’s network activation analysis.

For ProComReN modeling analysis, a signal network is obtained from a certain type of cell, the gene node data is Booleanized (or organized in “probability form”), and a Boolean network model is built to explore the dynamics of ASD synaptic disorders under the influence of different upstream signaling molecules. NIVaCaR and ProComReN are not sequential methods, but parallel. NIVaCaR identifies the activated subnetworks in mSiReN for each cell type and discovered the core module of EIF4E and EIF4EBP1 pair to further modeling in ProComReN. If these nodes in the activated subnetworks are used to construct a new network, it lacks coherence in the network structure.

##### The combinations of NIVaCaR and ProComReN

Next, We checked whether the result of subnetworks from NIVaCaR can be uniform to find a suitable and general network for further quantitative analysis by ProComReN.

- If NIVaCaR 1, ProComReN 1, then feasibility 0: Can’t find a uniform subnetwork for all cell types.
- If NIVaCaR 0, ProComReN 1, then feasibility 0: ProComReN can’t decide which jointed network to implement (E.G., 46 nodes are too many).
- If NIVaCaR 1, ProComReN 0, then feasibility 1: Lose the quantitative description of ProComReN.

For the structure of network to traing model, if we directly used the activated nodes and edges found directly by the NIVaCaR there could not have enough consistent node and edges Fig. 18.

##### The problem for core module-based inverse trace

But, if we use inverse trace Fig. 12 in the network, almost all nodes will be included after several neighbour incoming edges (finally discard five nodes). On the other hand, based on ”core module” to inverse trace network is also going to face the ”all incoming edges” problem, such as TSC2 has all the effect of other nodes, which any cell type can't have only. The third question is whether these edges direct from the interaction database exist in every cell type without using the RNA-seq data-based method, NIVaCaR.

##### The limit of union network to train model

We might need to use the unique or common network Fig. 19 and Fig. 20 from the results of all cell types and make further quantitative analyses by ProComReN. The conception of upstream nodes and the transmitter receptor is just used to NIVaCaR before the ProComReN pipeline. And in the process of ProComReN, we can focus more on customised subnetworks across cell types.

When employing the union network that incorporates all L or IN-type neurons, each node in the network becomes a fusion of multiple neuro-types. For instance, taking TSC2 as an example, it represents a blend of four different neuron types, with each type having its unique set of connections. Utilizing the ProComReN method to identify universal paths in the network based on specific cell type data may result in certain paths that are only relevant to that particular cell type. These paths might not exist in other cell types. Furthermore, the overall network structure, including the arrangement of edges, could vary significantly across different cell types.

In conclusion, the NIVaCaR and ProComReN methods are not congruent, making them unsuitable for direct combination. Both techniques aim to discover consistent pathways in a network that can match experimental data, but they differ in their underlying approaches. NIVaCaR relies on FC values, while ProComReN relies on gene expression data directly. Since the two methods are fundamentally different in their methodologies and data requirements, attempting to merge them directly may not yield meaningful or coherent results.

#### Supplementary Note 8 The Significance of mSiReN in Investigating Dysfunctions in Signal Transduction

##### Gene mutation, RNA, protein and phenotypic effects

The term "endophenotype" refers to a measurable and heritable characteristic or trait that is intermediate between a specific genetic variation and a complex phenotype or clinical manifestation of a disorder. Endophenotypes are believed to be more closely related to the underlying genetic mechanisms of a disorder than the observable symptoms or clinical diagnosis. They are often used in research to study the genetic and biological basis of complex traits, such as psychiatric disorders, by providing a more direct link to the underlying genetic factors. Endophenotypes can include physiological, biochemical, cognitive, neuroanatomical, or neurophysiological measures that are associated with a particular disorder. By studying endophenotypes, researchers aim to better understand the genetic basis and mechanisms of disorders and improve diagnostic accuracy and treatment approaches.

A gene mutation result in the production of a faulty RNA molecule, which can ultimately lead to the production of a dysfunctional protein. This can occur when the mutation alters the sequence of the gene in a way that disrupts the normal process of transcription, during which the DNA sequence is copied into RNA. The faulty RNA molecule can then be translated into a protein that is either incomplete or misfolded, which can result in the protein losing its normal function. In some cases, mutations in a single gene can result in downstream effects on other genes or proteins, which can lead to a convergence of phenotypic effects. Examples of such conditions include cystic fibrosis, sickle cell anemia, and Huntington's disease, all of which are caused by mutations in a single gene that result in the production of dysfunctional proteins with downstream effects on cellular processes.

##### The necessity of training quantitative model

Several reasons underscore the necessity of training quantitative models for RNA levels or transcriptomics:

1. RNA serves as a precursor for protein synthesis, and the RNA levels correlate positively with protein levels within a specific time frame.

2. ASD research differs from cancer research, where abundant cell lines and wet lab experiments can collect perturbation data to observe changes in signaling networks under various external stimuli.

3. While a significant portion of research focuses on gene regulatory networks (GRN), the impact of external stimuli on GRN networks and the downstream pathways of key genes remain uncertain.

4. ASD, a psychiatric disorder, involves the study of humans, as opposed to animal models, making it impossible to collect real-time samples for biochemical experiments to determine the molecular mechanisms in pathology.

##### The advantages of mSiReN in investigating dysfunction mechanisms

The mRNA Signaling-Regulatory Network (mSiReN) plays a crucial role in understanding the intricacies of signal transduction related to synapse plasticity and translation control 9. Here are the key advantages of mSiReN in investigating dysfunction mechanisms in the signal transduction network:

- mRNA as a Determinant of Protein Expression: mRNA serves as a pivotal factor influencing protein expression and provides valuable insights into potential dysfunctions in protein production within the transduction network. This extends beyond mere protein identification, encompassing regulatory interactions from the RNA world to the functional level of proteins. This understanding sheds light on the convergence of evidence associated with ASD, such as mRNA, non-coding RNA (ncRNA), and protein interactions in specific cell types, particularly in the DLPFC (Dorsolateral Prefrontal Cortex).

- Utilization of Abundant RNA Sequencing Data: mSiReN's modeling approach relies on extensive RNA sequencing expression data. This approach allows for the exploration of interactions between mRNA molecules, leading to a deeper understanding of the mechanisms underlying aberrations in the signal transduction network.

- Identification of Dysregulated RNA: mSiReN not only identifies dysregulated protein-coding RNA (mRNA) but also uncovers intricate regulatory relationships involving non-coding RNA. This is a significant contribution, as the protein-RNA interactions within the signal network, well-established in the ASD community, have not been fully explored. Our work introduces the possibility of aberrations at the RNA level, specifically RNA-ncRNA interactions, within the signal transduction network.

The comprehensive approach offered by mSiReN holds the potential to reveal novel insights into the molecular mechanisms contributing to ASD, especially concerning translation control in synaptic plasticity. Moreover, it opens avenues for the development of targeted therapeutic strategies aimed at specific endophenotypes of ASD.

### Supplementary Figure

| Group | ALL | INPV | INSST | INSV2C | INVIP | L23 | L4 | L56 | L56CC | Pathway |
|---|---|---|---|---|---|---|---|---|---|---|
| A | 35.3 | 3.7 | 3.5 | 5.8 | 7.1 | 6.1 | 8.3 | 2.9 | 4.1 | Neuronal System |
| | 30.6 | 4.8 | 4.6 | 7.5 | 7.3 | 6.8 | 5.7 | 2.7 | 5.4 | Transmission across Chemical Synapses |
| | 22.1 | 3.1 | 3.3 | 4.1 | 4.3 | 3.7 | 3.4 | <2.0 | 2.3 | Neurotransmitter receptors and postsynaptic signal transmission |
| | 6.2 | <2.0 | <2.0 | 4.5 | <2.0 | 4.3 | 3.3 | <2.0 | 4.5 | Neurotransmitter release cycle |
| | 11.3 | <2.0 | <2.0 | <2.0 | <2.0 | <2.0 | 3.8 | <2.0 | <2.0 | Protein-protein interactions at synapses |
| | 5.9 | <2.0 | <2.0 | <2.0 | <2.0 | <2.0 | <2.0 | <2.0 | <2.0 | Neurexins and neuroligins |
| | 4.0 | <2.0 | <2.0 | <2.0 | <2.0 | <2.0 | <2.0 | <2.0 | <2.0 | Synaptic adhesion-like molecules |
| B | 14.8 | <2.0 | 3.6 | <2.0 | 4.5 | 2.2 | 2.5 | <2.0 | 2.4 | Activation of NMDA receptors and postsynaptic events |
| | 8.5 | <2.0 | <2.0 | 3.8 | <2.0 | <2.0 | <2.0 | <2.0 | <2.0 | Glutamate binding, activation of AMPA receptors and synaptic plasticity |
| | 6.5 | 3.6 | <2.0 | <2.0 | <2.0 | 2.7 | <2.0 | <2.0 | <2.0 | GABA receptor activation |
| | 5.8 | <2.0 | 3.8 | <2.0 | <2.0 | <2.0 | <2.0 | <2.0 | <2.0 | Activation of kainate receptors upon glutamate binding |
| | 6.3 | <2.0 | <2.0 | 6.3 | <2.0 | <2.0 | <2.0 | <2.0 | <2.0 | GABA synthesis, release, reuptake and degradation |
| | 5.8 | <2.0 | <2.0 | <2.0 | <2.0 | 3.9 | <2.0 | <2.0 | 4.1 | Glutamate Neurotransmitter Release Cycle |
| | 4.1 | <2.0 | <2.0 | <2.0 | <2.0 | <2.0 | <2.0 | <2.0 | <2.0 | Acetylcholine Neurotransmitter Release Cycle |
| | 4.0 | <2.0 | <2.0 | <2.0 | <2.0 | <2.0 | <2.0 | <2.0 | <2.0 | Serotonin Neurotransmitter Release Cycle |
| | 4.0 | <2.0 | <2.0 | <2.0 | <2.0 | <2.0 | <2.0 | <2.0 | <2.0 | Norepinephrine Neurotransmitter Release Cycle |
| | 3.6 | <2.0 | <2.0 | <2.0 | <2.0 | <2.0 | <2.0 | <2.0 | <2.0 | Dopamine Neurotransmitter Release Cycle |
| C | 14.0 | <2.0 | 3.9 | <2.0 | 4.8 | 2.4 | 2.7 | <2.0 | 2.6 | Post NMDA receptor activation events |
| | 9.8 | <2.0 | 4.2 | <2.0 | <2.0 | <2.0 | 4.3 | <2.0 | <2.0 | Long-term potentiation |
| | 8.8 | <2.0 | 4 | <2.0 | <2.0 | <2.0 | <2.0 | <2.0 | <2.0 | CREB1 phosphorylation through NMDA receptor-mediated activation of RAS signaling |
| | 7.6 | <2.0 | 4.4 | <2.0 | <2.0 | <2.0 | <2.0 | <2.0 | <2.0 | Ras activation upon Ca2+ influx through NMDA receptor |
| | 6.4 | <2.0 | <2.0 | <2.0 | 4.6 | <2.0 | <2.0 | <2.0 | <2.0 | Activation of AMPK downstream of NMDARs |
| | 4.6 | <2.0 | <2.0 | <2.0 | <2.0 | <2.0 | <2.0 | <2.0 | <2.0 | CREB1 phosphorylation through the activation of CaMKII/CaMKK/CaMKIV cascasde |
| | 10.3 | <2.0 | <2.0 | <2.0 | <2.0 | <2.0 | <2.0 | <2.0 | <2.0 | Unblocking of NMDA receptors, glutamate binding and activation |
| | 7.2 | <2.0 | <2.0 | <2.0 | <2.0 | <2.0 | <2.0 | <2.0 | <2.0 | Assembly and cell surface presentation of NMDA receptors |
| | 6.1 | <2.0 | <2.0 | <2.0 | <2.0 | <2.0 | <2.0 | <2.0 | <2.0 | Negative regulation of NMDA receptor-mediated neuronal transmission |

NMDA and PostSynapse

Progressive enrichment analysis of DE genes in ASD

**Supplementary Figure 1. Synapse-related endophenotype as ASD dysfunction discovered from the evidence of Enrichment analyses of DEG in ASD.**

| ALL | INPV | INSST | INSV2C | INVIP | L23 | L4 | L56 | L56CC | Pathway |
|---|---|---|---|---|---|---|---|---|---|
| 15.2 | <2.0 | 3.1 | 2.9 | <2.0 | <2.0 | <2.0 | <2.0 | <2.0 | cAMP signaling pathway |
| 9.9 | <2.0 | <2.0 | 3.5 | <2.0 | <2.0 | <2.0 | <2.0 | <2.0 | cGMP-PKG signaling pathway |
| 8.9 | <2.0 | <2.0 | <2.0 | <2.0 | <2.0 | <2.0 | <2.0 | <2.0 | Rap1 signaling pathway |
| 8.3 | <2.0 | <2.0 | <2.0 | <2.0 | <2.0 | <2.0 | 2.2 | <2.0 | PI3K-Akt signaling pathway |
| 8.0 | <2.0 | <2.0 | <2.0 | <2.0 | <2.0 | <2.0 | <2.0 | <2.0 | Calcium signaling pathway |
| 7.9 | <2.0 | 2.9 | <2.0 | 3.9 | 2.4 | 2.1 | <2.0 | <2.0 | MAPK signaling pathway |
| 6.7 | 2.5 | 2.9 | <2.0 | <2.0 | <2.0 | <2.0 | <2.0 | <2.0 | Apelin signaling pathway |
| 6.5 | <2.0 | <2.0 | <2.0 | <2.0 | <2.0 | <2.0 | <2.0 | <2.0 | ErbB signaling pathway |
| 6.4 | <2.0 | <2.0 | <2.0 | <2.0 | <2.0 | <2.0 | <2.0 | 2.1 | HIF-1 signaling pathway |
| 4.3 | <2.0 | <2.0 | <2.0 | <2.0 | <2.0 | <2.0 | <2.0 | <2.0 | Hippo signaling pathway |
| 4.1 | <2.0 | <2.0 | <2.0 | <2.0 | <2.0 | <2.0 | <2.0 | <2.0 | Wnt signaling pathway |
| 3.9 | <2.0 | <2.0 | <2.0 | <2.0 | <2.0 | <2.0 | <2.0 | <2.0 | VEGF signaling pathway |
| 3.9 | <2.0 | 2 | <2.0 | <2.0 | <2.0 | <2.0 | <2.0 | <2.0 | Ras signaling pathway |
| 3.8 | <2.0 | <2.0 | <2.0 | <2.0 | <2.0 | <2.0 | <2.0 | <2.0 | mTOR signaling pathway |
| 3.7 | <2.0 | <2.0 | <2.0 | <2.0 | <2.0 | <2.0 | <2.0 | <2.0 | TNF signaling pathway |
| 3.5 | <2.0 | <2.0 | <2.0 | <2.0 | <2.0 | <2.0 | <2.0 | <2.0 | Phosphatidylinositol signaling system |
| 3.3 | <2.0 | <2.0 | <2.0 | <2.0 | <2.0 | <2.0 | <2.0 | <2.0 | Sphingolipid signaling pathway |

Color scale: 14, 12, 10, 8, 6, 4, 2, 0

**Supplementary Figure 2. KEGG enrichment of DE mRNA for Signal Transduction items in 8 types of brain cells.**

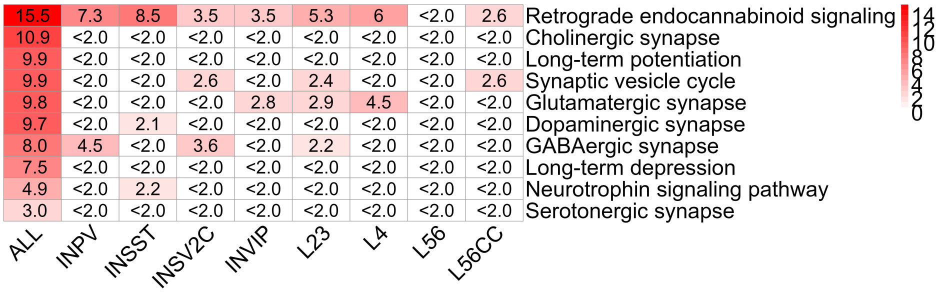

**Supplementary Figure 3. KEGG enrichment of DE mRNA for Neuronal System items in 8 types of brain cells.**

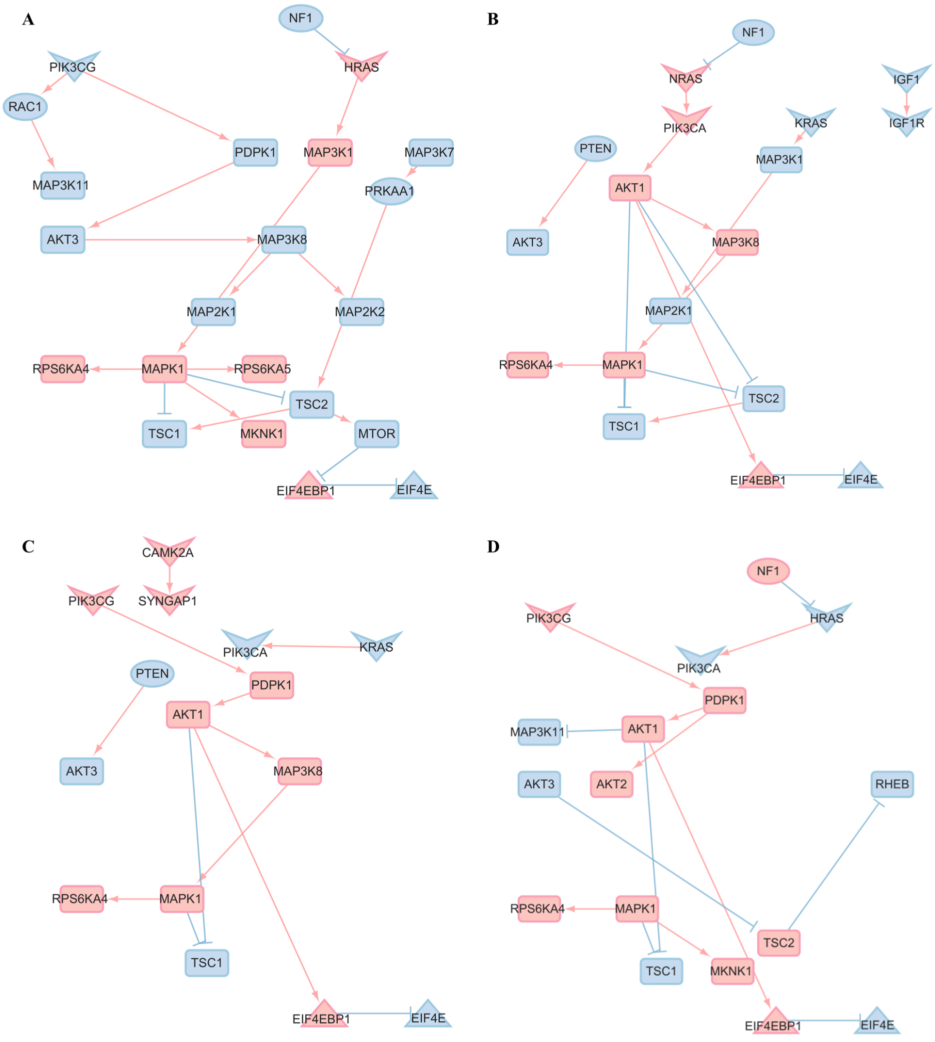

**Supplementary Figure 4. Employing NIVaCaR to identify a set of activated edges and nodes for L neurons within the mSiReN.**

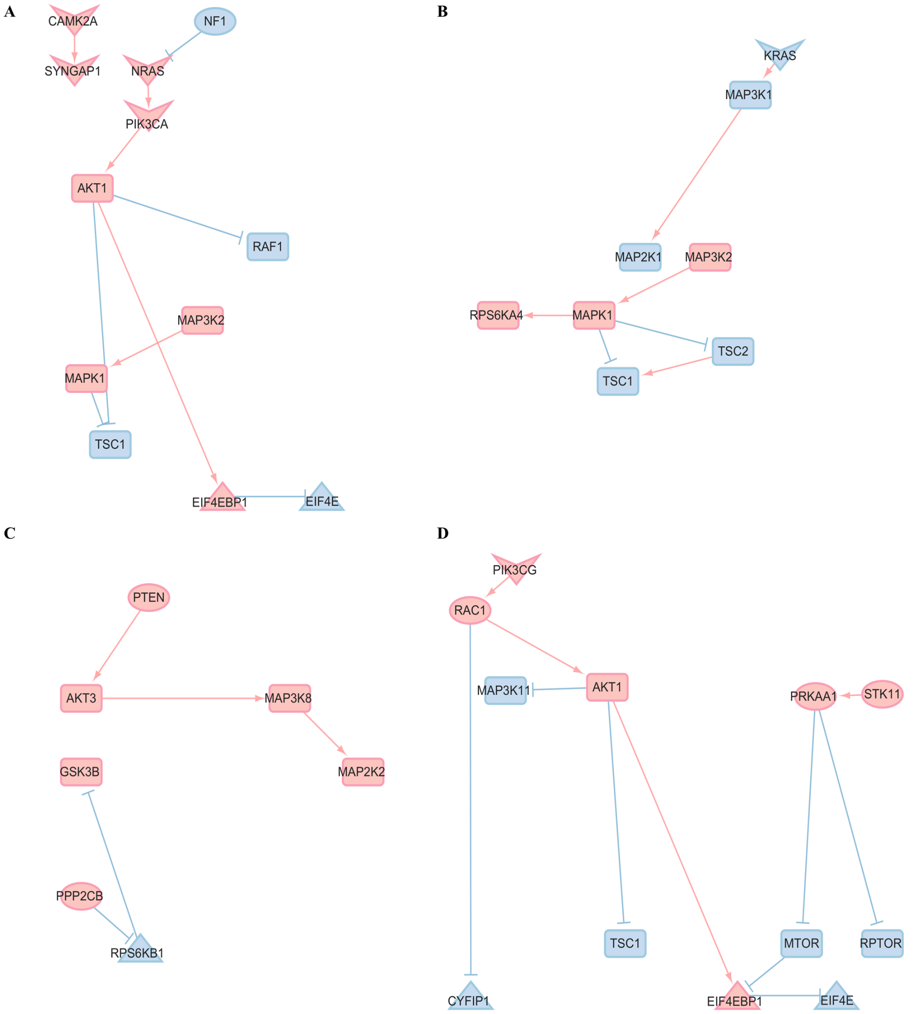

**Supplementary Figure 5. Employing NIVaCaR to identify a set of activated edges and nodes for IN neurons within the mSiReN.**

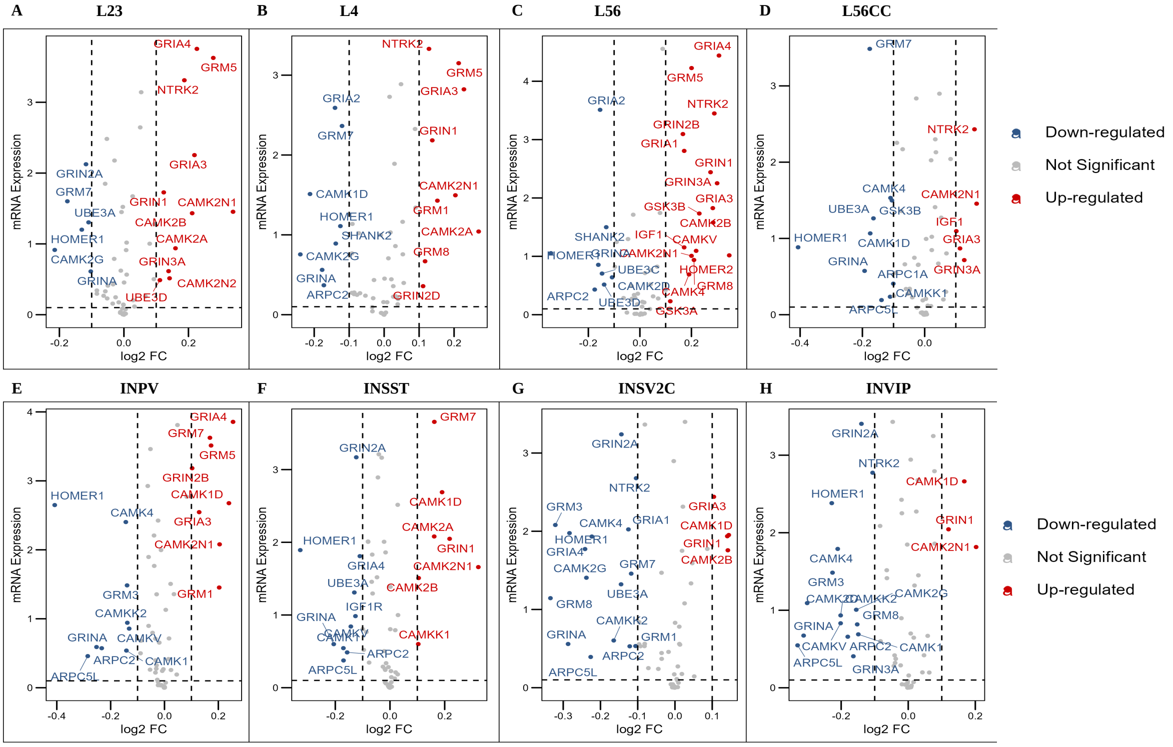

**Supplementary Figure 6. Volcanic map of expression and FC value for translation control related RNA in 8 cell types.**

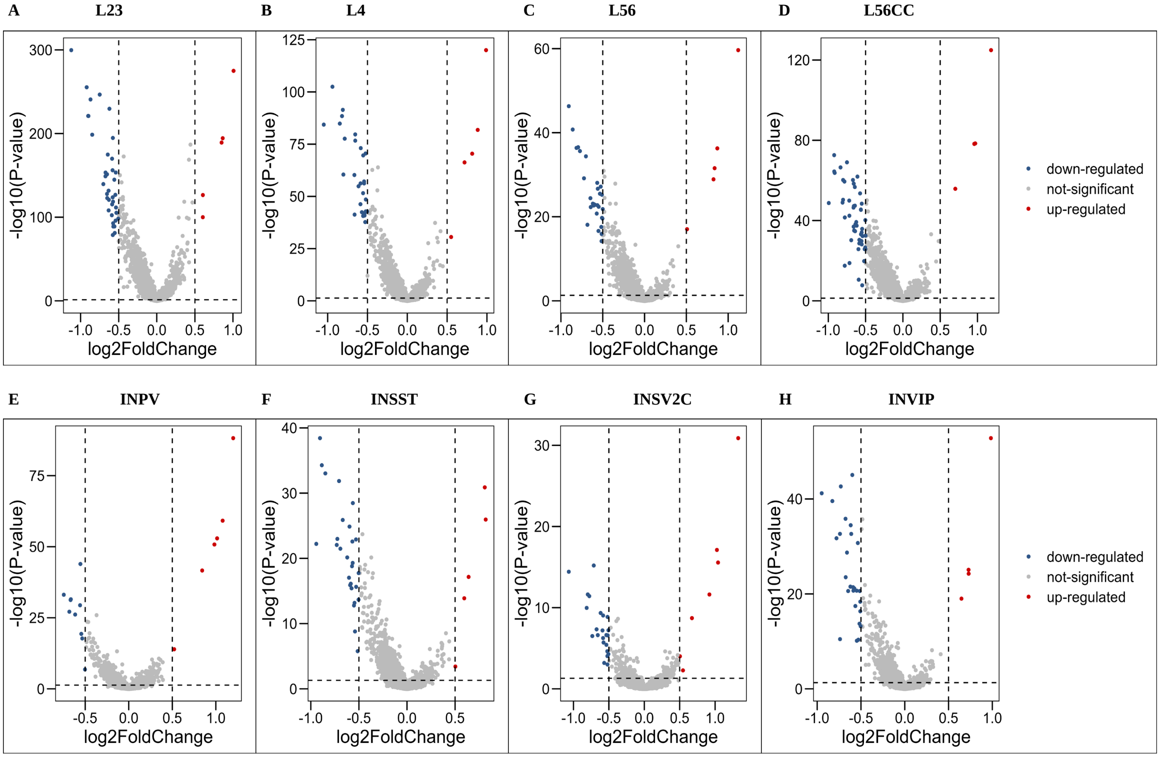

**Supplementary Figure 7. Volcanic map of DE P-Value and FC value for all DE RNAs in 8 cell types.**

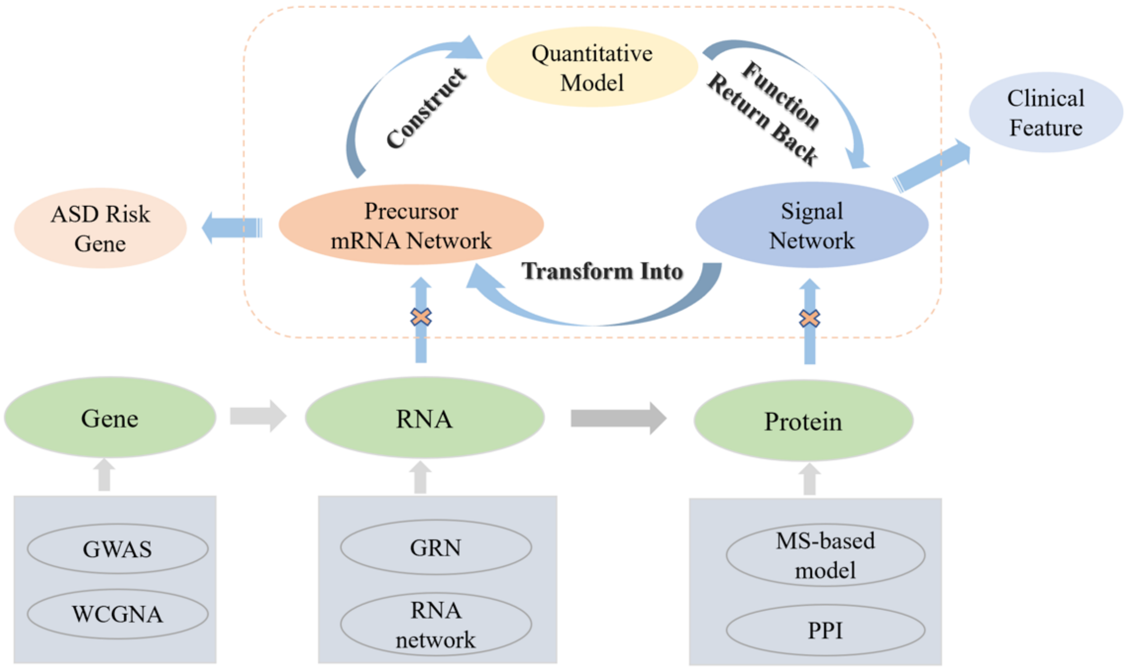

**Supplementary Figure 8. Methodology for Investigating Convergence in ASD Molecular Dysfunction.** The aim of our molecular convergence mechanism is to integrate studies at the individual molecular level. Our research combines the signaling network of protein regulation with transcriptional-level regulatory relationships in the context of the disease.

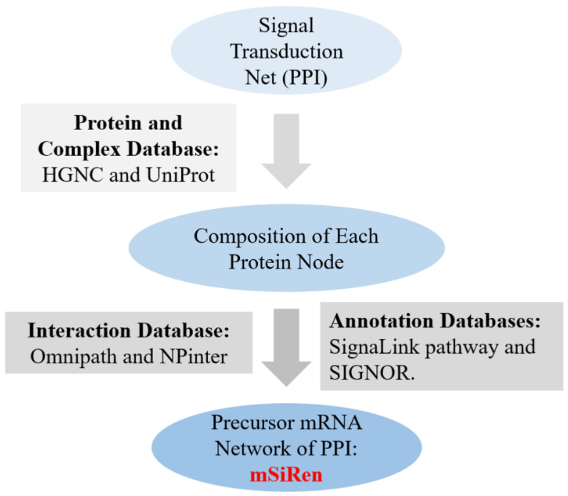

**Supplementary Figure 9. The construction process of mSiReN using databases.** We transformed the proteins in signal network into its corresponding precursor mRNAs based on databases HGNC and UniProt and found the relationship of regulation by interaction databases Omnipath and NPinter and annotation databases SignaLink pathway and SIGNOR.

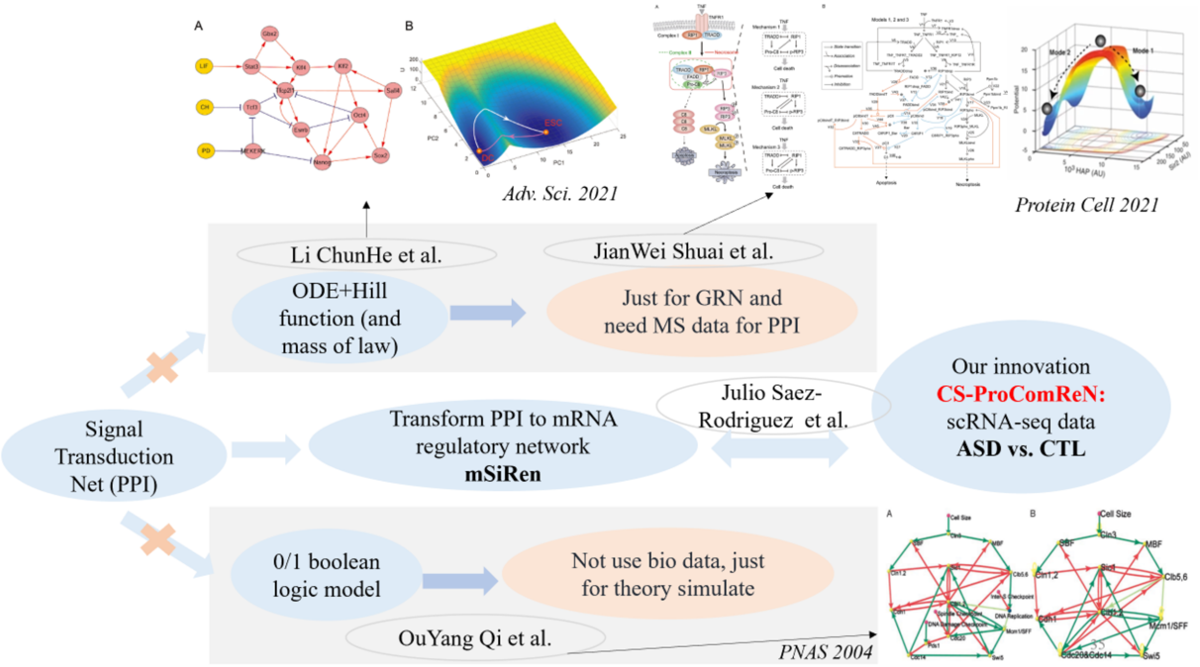

**Supplementary Figure 10. A comparison of the ProComReN method in the field of computational biology.** Instead of directly using ODE equations and pure mathematical modeling with a huge number of parameters, our ProComReN model integrates single-cell omics data and prior knowledge networks to investigate the molecular mechanisms of diseases.

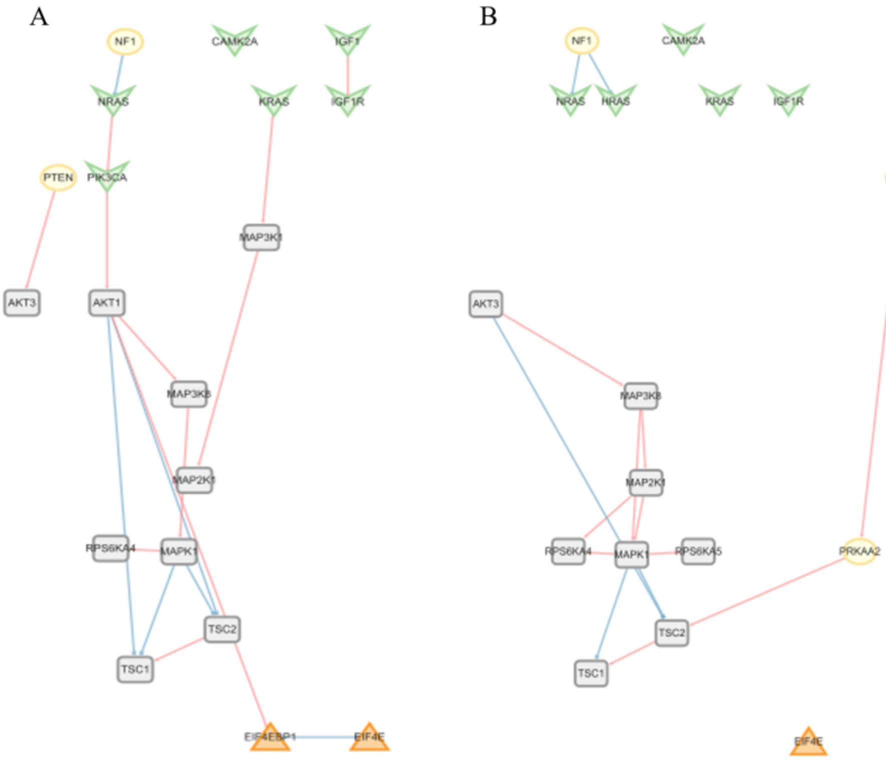

**Supplementary Figure 11. The results of NIVaCaR in L4 and the network of L4 DEGs.** A. The signal network is activated by the L4 by Carnival. B. Signal networks of L4 differentiated genes. The fold change is larger than 0.01 and the p-value is less than 0.05.

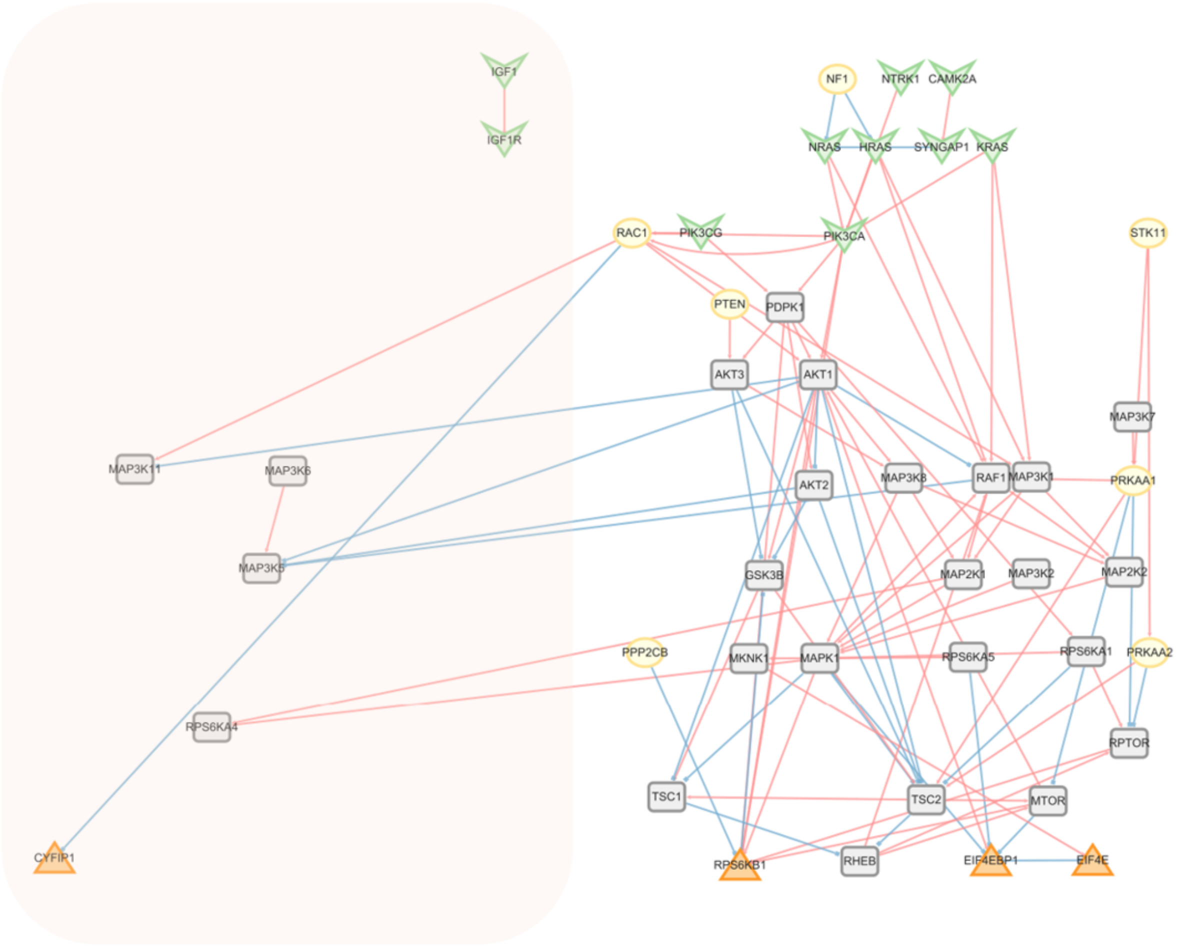

**Supplementary Figure 12. The mSiReN with core module-based inverse trace.** Delete some nodes in the pink area that are excluded from the inverse trace. It contains “IGF1”, “IGF1R”, “MAP3K11”, “MAP3K6”, “MAP3K5”, “RPS6KA4” and “CYFIP1”.

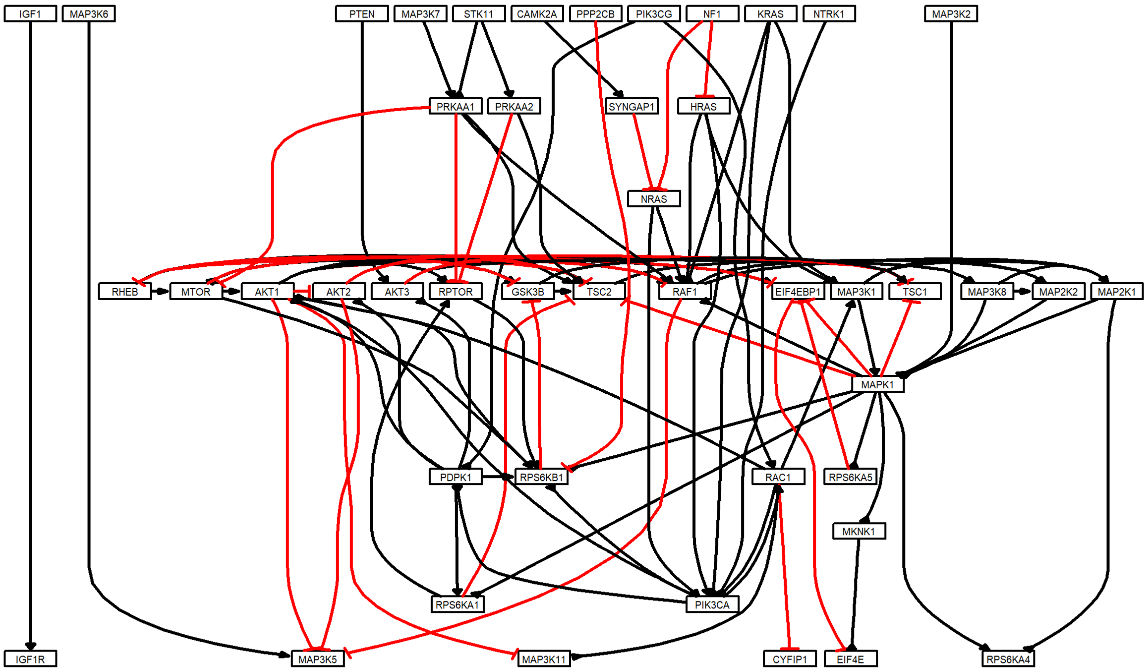

**Supplementary Figure 13. Based on the core module of 4EBP and EIF4E, we filtered some nodes from the original network which are excluded from the inverse trace.** Account: 7 nodes, 10 edges

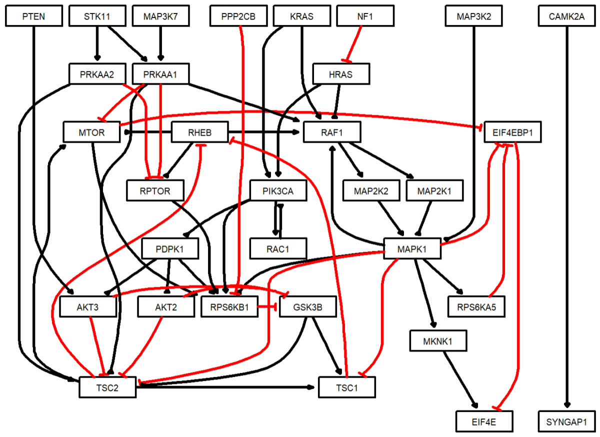

**Supplementary Figure 14. Delete some nodes lower than 20% expression in the cell type.**

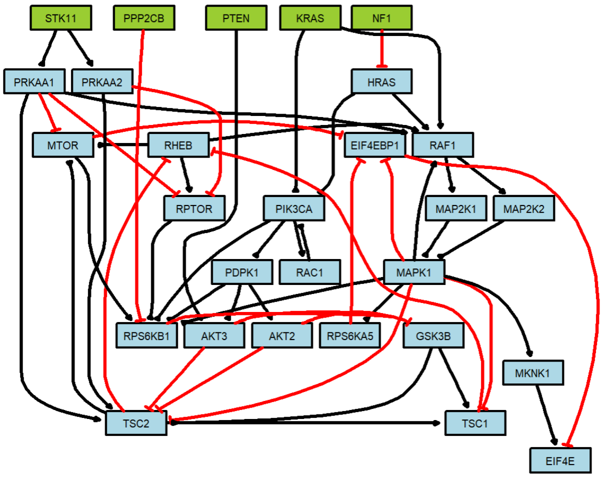

**Supplementary Figure 15. Delete upstream nodes based on topology.** Nodes: 28, edges: 53. Input nodes: KRAS, NF1, PPP2CB, PTEN, STK11.

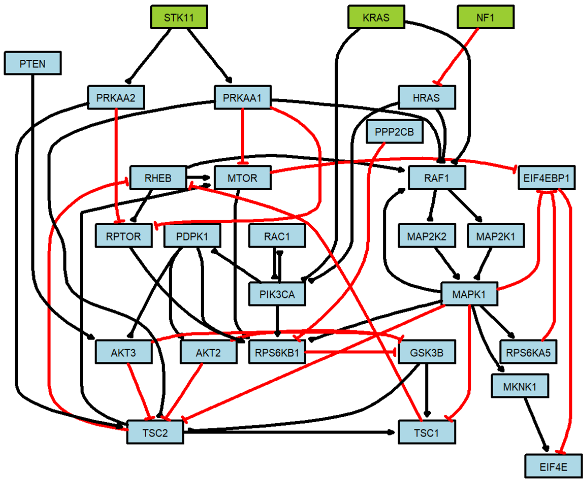

**Supplementary Figure 16. Narrow the network nodes from filtering homolog expression.**

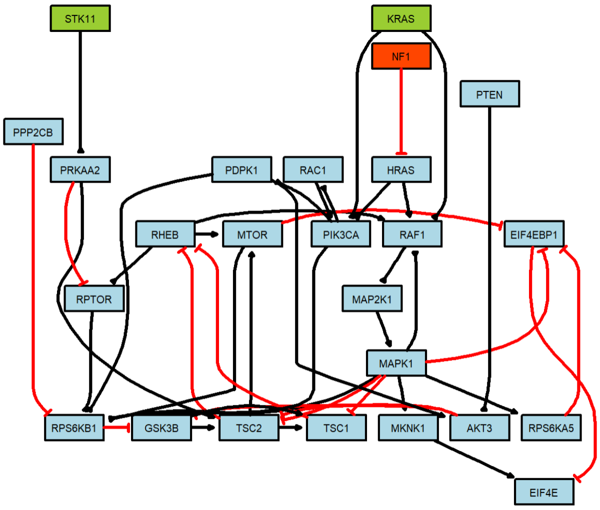

**Supplementary Figure 17. Final network after the reduced preprocess.**

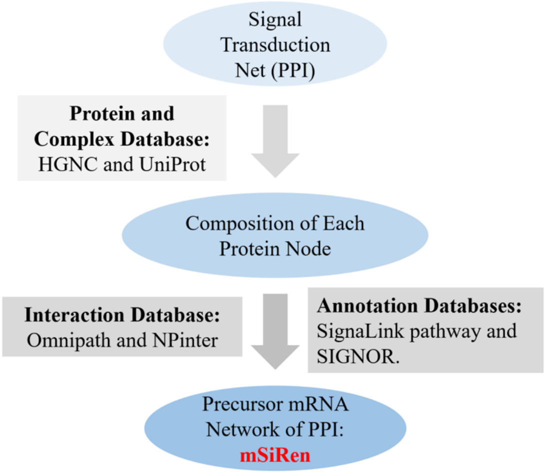

**Supplementary Figure 18. The subnetworks of L4 constructed from NIVaCaR directly.**

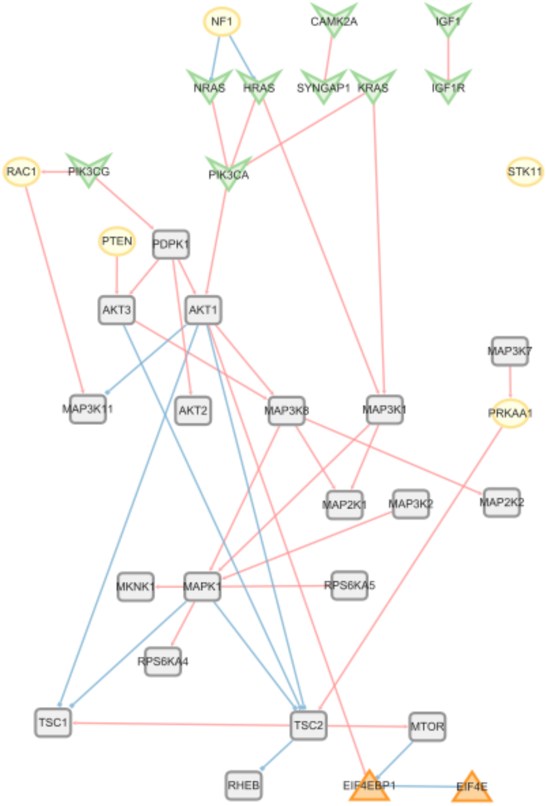

**Supplementary Figure 19. The Union network of L containing 35 nodes and 42 edges.**

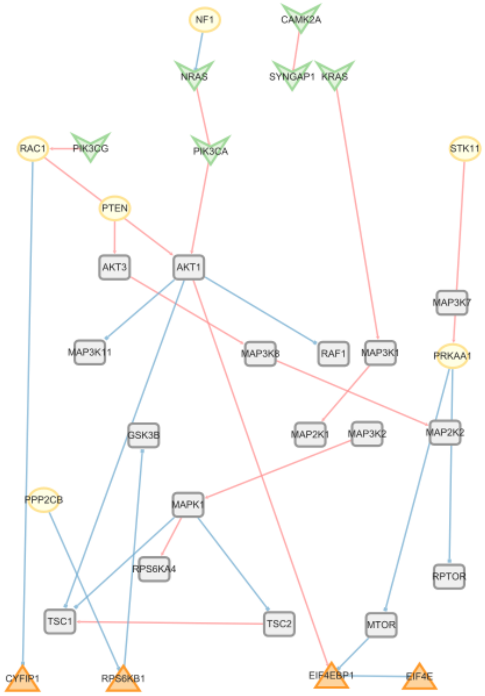

**Supplementary Figure 20. The Union network of IN containing 33 nodes and 28 edges.**

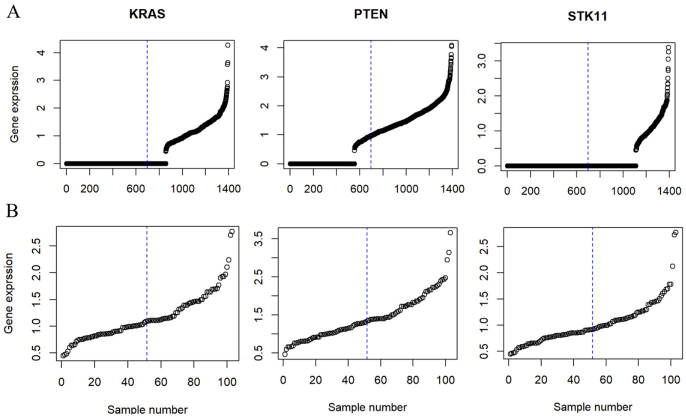

**Supplementary Figure 21. The number of cell in “L” cell types. A. Total “L” types cells sorted according to gene expression. B. All cells with no expression values were discard from (A).**

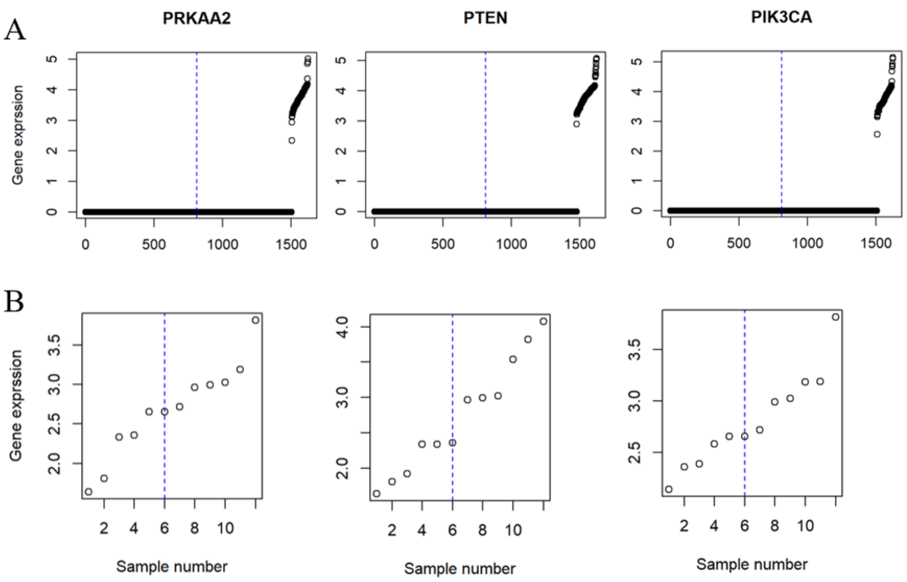

**Supplementary Figure 22. The number of cell in "NEU" cell types. A. Total "NEU" types cells sorted according to gene expression. B. All cells with no expression values were deleted from (A).**

### Supplementary Table

**Supplementary Table 1.** Cell number for different cell types in ASD and CTL from The high-throughout sc-RNA sequence data

| Cell Types | Num. total | Num. ASD | Num. CTL |
|---|---|---|---|
| AST-FB | 1571 | 1001 | 570 |
| AST-PP | 2395 | 1889 | 506 |
| Endothelial | 991 | 510 | 481 |
| IN-PV | 1481 | 829 | 652 |
| IN-SST | 1581 | 854 | 727 |
| IN-SV2C | 610 | 417 | 193 |
| IN-VIP | 1908 | 1289 | 619 |
| L2/3 | 5151 | 3146 | 2005 |
| L4 | 3195 | 2077 | 1118 |
| L5/6 | 1281 | 773 | 508 |
| L5/6-CC | 1562 | 920 | 642 |
| Microglia | 1027 | 635 | 392 |
| Neu-mat | 1459 | 945 | 514 |
| Neu-NRGN-I | 1164 | 616 | 548 |
| Neu-NRGN-II | 3024 | 1656 | 1368 |
| Oligodendrocytes | 3749 | 2055 | 1694 |
| OPC | 3207 | 1935 | 1272 |
| Total Num. | 35,356 | 21,547 | 13,809 |

**Supplementary Table 2.** The numbers of DE RNAs across different cell types from The high-throughout sc-RNA sequence data in ASD.

| Cell Types | Num. DE RNA |
|---|---|
| AST-FB | 145 |
| AST-PP | 172 |
| Endothelial | 172 |
| IN-PV | 56 |
| IN-SST | 89 |
| IN-SV2C | 96 |
| IN-VIP | 53 |
| L2/3 | 112 |
| L4 | 86 |
| L5/6 | 55 |
| L5/6-CC | 96 |
| Microglia | 167 |
| Neu-mat | 64 |
| Neu-NRGN-I | 122 |
| Neu-NRGN-II | 122 |
| Oligodendrocytes | 48 |
| OPC | 73 |

**Supplementary Table 3.** Network for source or end nodes in signal transduction network

| | **Label in signal transduction network (PPI and complex)** | **Precursor mRNA (gene)/ regulation** |
|---|---|---|
| Source: | "mGlu" | GRM (GRM1, GRM5) |
| | "iGlu" | GRIA1-4; GRIN1, GRIN2A-D, GRIN3A-B. |
| | "CAMKII" | CAMK2A |
| | "BDNF" | NTRK1, NTRK2, NTRK3 |
| | "SYNGAP1" | SYNGAP1 |
| | "IGF1" | IGF1, IGF1R |
| | "RAS" | HRAS, NRAS, KRAS4A and KRAS4B |
| | "PI3K" | PIK3CA_PIK3R1, PIK3CA/PIK3CB, PIK3CG |
| Target | "EIF-4E" | EIF4E |
| | "4E-BP" | EIF4EBP1, EIF4EBP1 |
| | "S6K1" | RKS6KB1, RKS6KB1, EIF4B/RPS6E |

**Supplementary Table 4.** Network process1 Filtering by signal annotation databases

| | 44 Initial nodes | Remain nodes |
|---|---|---|
| **Source Nodes** (Receptors and Upstream Signals) | CAMK1, CAMKID, CAMKIG, CAMK2A, CAMK2B, CAMK2D, CAMK2G, CAMK2N1, CAMK2N2, CAMK4, CAMKK1, CAMKK2, GRINIGRM2, GRIN2A, GRIN2B, GRIN2C, GRIN2D, GRIN3A, GRIN3B, GRM1, GRM3, GRM4, GRM5, GRM6, HRAS, IGF1, KRAS, GRM7, GRM8, IGFIR, NRAS, NTRKI, NTRK2, NTRK3, PIK3C2A, PIK3C2B, PIK3C2G, PIK3C3, PIK3CA, PIK3CB, PIK3CD, PIK3CG, SYNGAP1 | CAMK2A, HRAS, NTRK3, PIK3CA, IGEl, KRAS, IGEIR, SYNGAPI, PIK3CG, NRAS, NTRK1, NTRK2 |
| **Target Nodes** (Translation Control) | CYEIP1, CYFIP2, EIF4E, EIE4E1B, EIF4E2, EIE4E3, EIF4EBP1, EIF4EBP2, EIF4EBP3, RPS6KB1, RPS6KB2, RPS6KC1, RPS6KL1 | EIF4E, EIF4EBP1, RPS6KB1 |

**Supplementary Table 5.** Network process 2 Matching interaction database

| Nodes remain | Nodes discard |
|---|---|
| AKTI, AKT2, AKT3, CAMK2A, EIF4E, EIF4EBP1, GSK3B, HRAS, IGF1, IGFIR, KRAS, MAP2K1, MAP2K2, MAP3K1, MAP3K11, MAP3K2, MAP3K5, MAP3K6, MAP3K8, MAPK1, MKNKI, MTOR, NRAS, NTRKI, PDPKI, PIK3CA, PIK3CG, PTEN, RAFI, RHEB, RPS6KA1, RPS6KA4, RPS6KA5, RPS6KB1, RACI, RPTOR, SYNGAP1, TSCl, TSC2 | MAP3K10, NTRK2, MAP3K12, MAP3K13, NTRK3, RAC2, MAP3K20, MAP3K3, STK11, MAP3K4, MAP3K7, MAP3K9 |

**Supplementary Table 6.** Network process 3 Adding some super nodes about synapese plasticity

| Super nodes supplement | Final discard super nodes |
|---|---|
| ADNP, PP2A, CYFIP1, AMPK, EN2, LKB1 (STK11), FMRP, NE1 | ADNP, NMDAR(GRIN1), EN2, NMDAR,(GRIN2A-D), FMRP, mGluR1/5 (GRM1,GRM5) |

**Supplementary Table 7.** The NIVaCaR variables and their descriptions

| **Variable** | **Description** |
|---|---|
| $c_{j,k} \in \{-1, 0, 1\}$ | Activation/inhibition state of measured species $j$ for cell type $k$ |
| $x_{j,k} \in \{-1, 0, 1\}$ | Predicted activation/inhibition state of species $j$ for cell type $k$ |
| $x^+_{j,k} \in \{0, 1\}$ | Potential of node $j$ to be activated for cell type $k$ |
| $x^-_{j,k} \in \{0, 1\}$ | Potential of node $j$ to be inhibited for cell type $k$ |
| $u^+_{i,k} \in \{0, 1\}$ | Potential of interaction $i$ to activate its target node for cell type $k$ |
| $u^-_{i,k} \in \{0, 1\}$ | Potential of interaction $i$ to inhibit its target node for cell type $k$ |
| $\sigma_{i,k} \in \{-1, 1\}$ | Sign of interaction $i$ for cell type $k$ |
| $d_{j,k} \in \{0, M\}$ | Auxiliary distance variables assigned to each node $j$ where $M$ is a sufficiently large number (default: $M = 100$) for cell type $k$ |
| $A_{j,k} \in \{0, 2\}$ | Auxiliary variable representing the absolute difference between the inferred and measured species $j$ for cell type $k$ |

**Supplementary Table 8.** Nodes filtering from homolog.

| Ortholog Symbol HGNC/ Subunit | Node Label in Network | AveExpr |
|---|---|---|
| AKT2 | AKT (PKB) | 0.3607864 |
| AKT3 | AKT (PKB) | 2.90043385 |
| MAP2K1 | MEK1/2(MAP2K) | 1.00998264 |
| MAP2K2 | MEK1/2(MAP2K) | 0.29219968 |
| PRKAA1 | AMPK | 0.26095921 |
| PRKAA2 | AMPK | 1.11882331 |

**Supplementary Table 9.** Different types of biological interactions modelled by different Boolean functions and their algebraic representations.

| Biological equivalent | Graphical form | Algebraic computation |
|---|---|---|
| Activation | $A \rightarrow Z(k)$ | $Z_{t+1} = A_t * k$ |
| Inhibition | $A \dashv Z(k)$ | $Z_{t+1} = 1 - A_t * k$ |
| Complex formation | $A \textit{ and } B \rightarrow Z(k)$ | $Z_{t+1} = A_t * B_t * k$ |
| Competitive interaction | $A \textit{ or } B \rightarrow Z(k)$ | $Z_{t+1} = 1 - [(1 - A_t) * (1 - B_t) * k]$ |
| Non-competitive interaction | $A \rightarrow Z(k_1)B \rightarrow Z(k_2)$ | $Z_{t+1} = A_t * k_1 + B_t * k_2(k_1 + k_2 = 1)$ |
